# Human IL-10-producing B cells have diverse states induced from multiple B cell subsets

**DOI:** 10.1101/2021.09.01.458645

**Authors:** Marla C. Glass, David R. Glass, John Paul Oliveria, Berenice Mbiribindi, Carlos O. Esquivel, Sheri M. Krams, Sean C. Bendall, Olivia M. Martinez

## Abstract

Regulatory B cells (Bregs) can suppress immune responses through the secretion of IL-10 and other anti-inflammatory cytokines. This immunomodulatory capacity holds therapeutic potential, yet a definitional immunophenotype for enumeration and prospective isolation of B cells capable of IL-10 production remains elusive. We therefore applied mass cytometry to simultaneously quantify cytokine production and immunophenotype in human peripheral B cells across a range of stimulatory conditions and timepoints. While multiple B cell subsets produced IL-10, no phenotype uniquely identified IL-10^+^ B cells and a significant portion of IL-10^+^ B cells co-expressed the proinflammatory cytokines IL-6 and TNFα. Despite this heterogeneity, we found operationally tolerant liver transplant recipients had a unique enrichment of IL-10^+^, but not TNFα^+^ or IL-6^+^, B cells as compared to transplant recipients receiving immunosuppression. Thus, human IL-10-producing B cells constitute an induced, transient state arising from a diversity of B cell subsets that may contribute to maintenance of immune homeostasis.

## Introduction

In addition to their established roles in antibody production and antigen presentation, B cells support homeostasis of the immune system and regulation of inflammation. B cells with immunosuppressive properties, defined by expression of the hallmark immunoregulatory cytokine, IL-10, have been termed ‘regulatory B cells’ (Bregs) (Fillatreau et al. 2002; Mizoguchi et al. 2002; Mauri et al. 2003; Iwata et al. 2011). Granzyme B (Zhu et al. 2017), IL-35 (Shen et al. 2014; Wang et al. 2014), and other secreted molecules (Lee et al. 2014; Parekh et al. 2003), as well as surface receptors that alter cellular function (Catalán et al. 2021) have also been implicated in Breg immunomodulatory activity. Identification of an IL-10-producing B cell population with immunomodulatory activity was first reported in mice, where it was demonstrated that IL-10^+^ B cells were critical for prevention or control of autoimmune disease and chronic inflammation (Wolf et al. 1996). A myriad of phenotypes and regulatory functions of Bregs have since been established using murine models of disease (Ran et al. 2020; Jansen et al. 2021), yet the relationship and relevance of murine IL-10^+^ B cells to the human immune system requires further study (Lighaam et al. 2018; Rosser et al. 2014; Baba et al. 2020).

Human IL-10^+^ B cells have been implicated in autoimmunity (Iwata et al. 2011; Matsumoto et al. 2014; Flores-Borja et al. 2013), alloimmunity (Nova-Lamperti et al. 2016; Chesneau et al. 2014; Cherukuri et al. 2021; Newell et al. 2015; Silva et al. 2012), allergy (van de Veen et al. 2013; Oliveria et al. 2018), cancer (Michaud et al. 2021; Wang et al. 2015; Mehdipour et al. 2019; Budczies et al. 2021) and infection (Das et al. 2012; Lopez-Abente et al. 2018). A wide variety of Breg cell surface phenotypes were described in these studies, some of which are distinct from established murine Breg phenotypes (Jansen et al. 2021). Thus, a significant barrier in understanding and defining human Bregs has been the lack of an encompassing immunophenotypic signature to identify cells with IL-10 production capacity. As in the mouse, human Bregs are functionally defined by IL-10 expression and have been primarily studied in disease states characterized by immune alterations. Along these lines, CD24^hi^CD38^hi^ transitional B cells, CD9^+^ B cells, CD24^hi^CD27^+^ memory B cells and CD27^int^CD38^+^ plasmablasts were all established to produce IL-10, and in some cases to possess IL-10-dependent regulatory properties in healthy humans, yet were diminished or absent in human disease states (Iwata et al. 2011; Matsumoto et al. 2014; Menon et al. 2016; Brosseau et al. 2018; Blair et al. 2010). Collectively, these findings suggest that human IL-10-producing Bregs participate in maintenance and regulation of immune homeostasis. However, the dynamics of immunoregulatory IL-10 versus pro-inflammatory cytokine production by B cell functional subsets across healthy individuals and the impact of different activating stimuli remains to be verified by a systemic approach. Furthermore, the breadth of IL-10-producing B cell immunophenotypes is not well-described outside of specific activating contexts or disease states.

In clinical transplantation, the term operational tolerance describes a state whereby organ transplant recipients maintain stable graft function over an extended period in the absence of immunosuppression (IS). It remains unclear if this state of operational tolerance involves an active immunoregulatory process. Nevertheless, spontaneously tolerant renal transplant recipients demonstrated an increase in total B cells, particularly CD38^+^CD24^+^IgD^+^ transitional and naïve subsets, as well as an upregulated expression of B cell–associated genes by peripheral blood mononuclear cells (PBMCs), relative to their immunosuppressed counterparts (Newell et al. 2015; Pallier et al. 2010). *Chesneau et al* further demonstrated that operationally tolerant kidney recipients had increased proportions of CD20^+^CD24^hi^CD38^hi^ transitional B cells and CD20^+^CD38^lo^CD24^lo^ naïve B cells, and increased IL-10 production from activated CD19^+^ B cells as compared to healthy subjects and stable transplant recipients (Chesneau et al. 2014). Functionally, human IL-10^+^ B cells from healthy individuals and stable immunosuppressed kidney transplant recipients have been shown to decrease effector CD4^+^ T cell proliferation and type I proinflammatory cytokine or type II interferon production *in vitro* (Nova-Lamperti et al. 2016; Cherukuri et al. 2014). Additionally, Granzyme B^+^ (GzmB^+^) B cells from stable immunosuppressed and operationally tolerant renal transplant recipients also inhibited CD4^+^ T cell proliferation (Chesneau et al. 2015). Notably, the role of Bregs in transplantation has focused extensively on kidney allograft recipients, with limited published data on the contribution of these cells to operational tolerance of other transplanted organs. Comparison of peripheral blood transcriptional profiles in operationally tolerant kidney and liver recipients indicated a unique enrichment and expansion of B cell-associated genes in tolerant kidney recipients (Lozano et al. 2011). Thus, the relevance of B cells to tolerance in non-kidney transplant recipients remains unknown.

In homeostasis, B cells express only low or transient levels of IL-10, which obligates the use of *ex vivo* stimulation to clearly distinguish which B cells have IL-10-producing capacity. Moreover, a wide variety of stimuli have been reported for elucidating this across Breg studies (Iwata et al. 2011; Tedder 2015; Mauri and Bosma 2012; Baba et al. 2015; Mauri and Menon 2017). These studies have, however illuminated the importance of Toll-like receptor (TLR) engagement and CD40 activation in potent IL-10 induction (Iwata et al. 2011; Lighaam et al. 2018; Nova-Lamperti et al. 2016; van de Veen et al. 2013) and the augmentation of IL-10 expression by exogenous cytokines, such as IL-2, IL-35 and IL-21 (Tedder and Leonard 2014; Yoshizaki et al. 2012; Dambuza et al. 2017). The role of cognate stimulation via the B cell receptor (BCR) is less clear and, in some cases, has been shown to interfere with IL-10 production (Iwata et al. 2011; Parekh et al. 2003; Nova-Lamperti et al. 2016; Yanaba et al. 2009; Bankó et al. 2017; Maseda et al. 2012).

While there is likely an impactful role of Bregs in both human disease and immune tolerance, their presence, origins, cell states and variation across healthy individuals remains elusive. This is partially owed to the low level of IL-10 expression by B cell subsets *in vivo* (Fillatreau et al. 2002; Yanaba et al. 2009; Mauri and Menon 2017). Furthermore, Bregs have been frequently defined in the context of a specific disease or animal model of disease, through a single mode of *ex vivo* stimulation and without consideration of the substantial changes in B cell surface phenotype that occur upon potent activation. Collectively, these caveats have led to conflicting reports on what Bregs are and how they function.

Previously, we defined the spectrum of human B cell subsets across multiple tissues and related their phenotypic, metabolic, and immune signaling profiles (Glass et al. 2020). Here, we focus specifically on human Breg cells and apply mass cytometry to achieve high-dimensional characterization of IL-10^+^ B cells, with simultaneous quantification of 38 cell identity and state defining molecules, from healthy individuals and operationally tolerant liver allograft recipients. We evaluate the impact of TLR ligands, CD40 stimulation and exogenous cytokines on the generation of IL-10^+^ B cells and determine the potential for discrete B cell subsets to give rise to IL-10^+^ populations. Our results challenge the paradigm that Bregs constitute a distinct, homogenous B cell subset that is identifiable by a single phenotypic profile. Instead, using prospective isolation and a live cell tracing assay, our findings indicate that IL-10^+^ B cells can emerge from numerous previously defined B cell subsets. Moreover, the predominate IL-10^+^ B cell subset may vary depending on the stimuli and the immune status of the individual. Our study highlights the diversity of IL-10^+^ B cell states and suggests phenotypic heterogeneity within activated B cell compartments may help maintain a balance between proinflammatory and immunoregulatory responses.

## Results

### Emergence of human IL-10-producing B cells depends on cytokine and activation environment

Comparisons of human IL-10^+^ CD19^+^ B cells across studies have been challenging because a variety of *ex vivo* stimulation approaches have been used, and often the B cells were isolated from individuals with immune-related disease. Nonetheless, previous studies cooperatively indicate TLR and CD40 signaling, and cytokines such as IL-21 and IL-35, are important in generation of IL-10-producing B cells (Iwata et al. 2011; Tedder 2015; Mauri and Bosma 2012; Baba et al. 2015; Mauri and Menon 2017). To characterize and compare the effects of different B cell stimuli on IL-10 induction and B cell phenotype, we activated human PBMCs from healthy individuals for 72 hours with TLR ligands, anti-CD40, and exogenous cytokines, individually and in combination. We then applied mass cytometry to assess the expression of 24 surface and 14 intracellular proteins on total CD19^+^ B cells (Figure 1A). We first compared the various stimulatory conditions (Figure 1B) for the ability to elicit IL-10 production from B cells in PBMC from three healthy individuals (Figure 1C). TLR7/8 (R848) and TLR9 (CpG) stimulation, either alone or in combination with CD40 activation, yielded higher proportions of IL-10-producing B cells than did TLR4-based (LPS) stimulation (Figure 1D). Of the 11 stimulation conditions tested, the cocktail of CpG, anti-CD40, IL-2, IL-21, and IL-35 (stimulation 10) was most effective in eliciting IL-10 production from B cells of healthy individuals (mean of 34.1% IL-10^+^ B cells) (Figure 1D).

**Figure 1:**
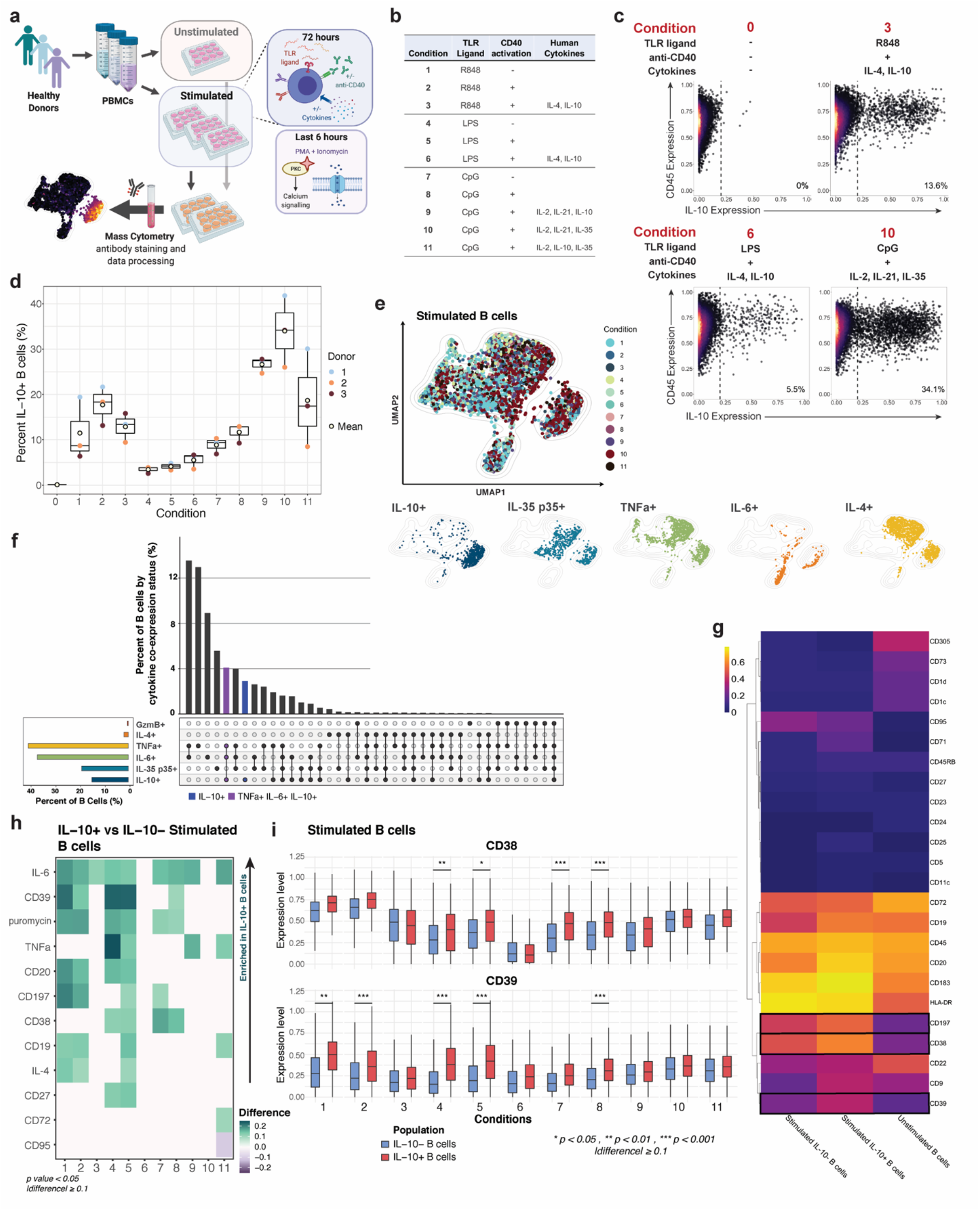
IL-10^+^ B cell immunophenotype and cytokine production varies by activation microenvironment. **(A)** Experimental workflow **(B)** Summary of *in vitro* stimulatory conditions. **(C)** Exemplative biaxial plots demonstrating differential IL-10 expression by B cells from the same pooled donors under varying stimulation conditions. Unstimulated PBMCs are depicted as condition 0. **(D)** Percent IL-10^+^ B cells by condition and by individual (colored dots). Unstimulated B cell data is depicted as condition 0. Line depicts median, box depicts interquartile range (IQR), and whiskers depict IQR+-1.5*IQR. **(E)** UMAP projection of donor-pooled B cells colored by stimulatory condition (upper panel) and by positive status of the indicated cytokine (lower panels). Mass cytometry data was subsampled equally by condition. Manifold was derived using only expression of phenotypic markers. **(F)** Upper bar graph indicates the percent of stimulated B cells (upper bar graph) by cytokine (IL-4, TNF*a*, IL-6, IL-35p35, IL-10) and GzmB expression status, as determined by Boolean gating of cytokine-positive cells. Cytokine and GzmB co-expression status of populations are defined in the lower dot plot panel. Left lower bar graph depicts percent of cytokine/GzmB-positive B cells regardless of co-expression, across all stimulation conditions. Data was subsampled equally by condition. **(G)** Median surface marker expression for unstimulated total B cells and IL-10^+^ and IL-10^-^ stimulated B cell subsets. Boxes indicate non-lineage markers with significant upregulation in IL-10^+^ B cells as compared to IL-10^-^ B cells in 4 or more (of 11) stimulatory conditions. **(H)** Relative difference in median expression of surface markers and cytokines in IL-10^+^ versus IL-10^-^ stimulated B cells by activating condition (1-11). Expression only shown for markers and cytokines with difference greater than or equal to 0.1 and a p value less than 0.05. P values denote result of Kolmogorov-Smirnov test with Bonferroni multiple hypothesis correcting. **(I)** Expression levels of CD38 and CD39 for IL-10^-^ (blue) versus IL-10^+^ (red) B cells by stimulation condition. P values denote result of Kolmogorov-Smirnov test with Bonferroni multiple hypothesis correcting. Line depicts median, box depicts interquartile range (IQR), and whiskers depict IQR+-1.5*IQR. * p < 0.05, ** p < 0.01, and *** p < 0.001.

We applied UMAP (Becht et al. 2019) to visualize the range of phenotypes and cytokine profiles induced by the 11 different stimulation conditions (Figure 1E, upper panel). IL-10^+^ cells were largely, though not exclusively, located in one island of the UMAP projection but IL-35^+^ (p35), TNFα^+^, IL-6^+^ and IL-4^+^ B cells also occupied the same island indicating the polyfunctionality of IL-10^+^ B cells (Fig 1E, lower panel). To quantify this polyfunctionality, we equally subsampled total B cells by stimulatory condition *in silico* and determined the proportion of those B cells that were able to produce IL-10 alone or in combination with IL-4, TNFα, IL-6, and/or the cytotoxic mediator GzmB (Figure 1F). While B cells that were exclusively IL-10 producers could be detected (blue bar), the greatest proportion of IL-10^+^ B cells also produced TNFa and IL-6 (purple bar) with lesser proportions of IL-10^+^ B cells also producing IL-4, GzmB, and the p35 subunit of IL-35. This profile held true regardless of stimulatory condition (Figure S1).

To identify features that differentiate IL-10^+^ B cells from IL-10^-^ B cells across the 11 different stimulation conditions, we compared expression of 24 surface markers and four cytokines between the two subsets (Figure 1G, H). IL-10^+^ B cells had significantly higher expression levels of IL-6 and/or TNFα as compared to IL-10^-^ B cells in most stimulatory conditions (p<0.05, Kolmogorov-Smirnov (KS) test with Bonferroni correction). IL-10 and TNFα co-induction in *ex vivo* stimulated human B cells is consistent with reports from other groups (Lighaam et al. 2018; Cherukuri et al. 2014, 2021). Co-expression of IL-10 and IL-4 was significantly elevated with TLR4 and CD40 stimulation and with TLR7/8 ± CD40 stimulation (conditions 1, 2 and 5) (Figure 1H). In all stimulatory conditions, IL-10^+^ B cells show a highly activated B cell phenotype (Figure 1H, S1), which included upregulation of HLA-DR, CD38, CD183 (CXCR3), and CD197 (CCR7) and increased protein synthesis as indicated by puromycin labeling (Kimmey et al. 2019). While the median expression of these molecules was generally higher in IL-10^+^ B cells than IL-10^-^ cells (Figure 1H), IL-10^+^ and IL-10^-^ stimulated B cells had similar expression profiles (Figure 1G).

Amongst individual surface markers, CD39, an immunomodulatory ectonucleotidase, most frequently demonstrated significant association with IL-10 expression by B cells (within conditions 1, 2, 4, 5 and 8) and generally had elevated median expression in IL-10^+^ stimulated B cells as compared to IL-10^-^ stimulated B cells (Figure 1H, I). CD38, an activation molecule and marker of rapidly proliferating cells, was the second most frequent non-lineage surface molecule to be significantly upregulated in IL-10^+^ B cells within the screen of stimulatory conditions. Notably, condition 11 induced a unique IL-10^+^ B cell population with significantly upregulated CD72, a BCR co-receptor with regulatory activity, and downregulated CD95, a memory-associated marker (Figure 1H). CD9 was also upregulated in IL-10^+^ stimulated B cells as compared to IL-10^-^ B cells, in all but one stimulatory condition (9), but this expression change was not significant (Figure 1G, Figure S2). We also noted distinct CD23 upregulation and greater induction of CD23^+^ IL-10^+^ B cells upon R848- or LPS-based stimulation with CD40 activation and exogenous IL-4 and IL-10, as compared to all other conditions (Figure S1). Interestingly, CD23 is a negative regulator of BCR signaling as well as IgG/IgE antibody responses. Collectively, we observed that IL-10^+^ B cell surface phenotypes and cytokine profiles vary according to the specific stimulation environment. While CD39 most significantly identified exclusive IL-10-producing cells, no single phenotype captured these populations across all modes of activation.

### Pro-inflammatory cytokines expression precedes B cell IL-10 induction

Prior studies indicate that the peak of IL-10 production as well as proinflammatory cytokine co-expression by B cells can vary significantly depending upon the method and duration of *ex vivo* stimulation (Iwata et al. 2011). In our stimulation screen, the culture of PBMCs in the presence of CpG, anti-CD40 and exogenous recombinant human (rh) IL-2, IL-21, and IL-35 (condition 10) yielded maximal IL-10 expression by peripheral B cells. To map the temporal dynamics of cytokine production and marker expression by B cells, we recovered total PBMCs after 6, 12, 24, 48, 60 and 72 hours of stimulation and applied mass cytometry and high-dimensional analysis, to evaluate B cell expression profiles (Figure 2A). The frequency of IL-10^+^ B cells peaked at 48 hours of *ex vivo* stimulation and this observation was consistent across individuals (Figure 2B). Notably, high expression of the pro-inflammatory cytokines, TNFα and IL-6, preceded IL-10 induction (Figure 2C-D, S3-4). At the earliest time point analyzed (6 hours), most B cells expressed both TNFα and IL-6, while <1% of cells expressed IL-10 (Figure 2C, S4). TNFα^+^IL-10^+^ polyfunctional B cells began to emerge at 12-24 hours and by 48 hours post-stimulation, a significant population of B cells that expressed only IL-10 was present, while the TNFα^+^ B cell population substantially contracted (Figure 2C-D, S3). The TNFα and IL-10 co-expression profile of total B cells remained largely stable between 48 to 72 hours post-stimulation. These data suggest that in the early phase following activation, B cells predominantly provide inflammatory signals that are subsequently replaced by a more regulatory B cell state, including a peak of IL-10 production at 48 hours of stimulation.

**Figure 2:**
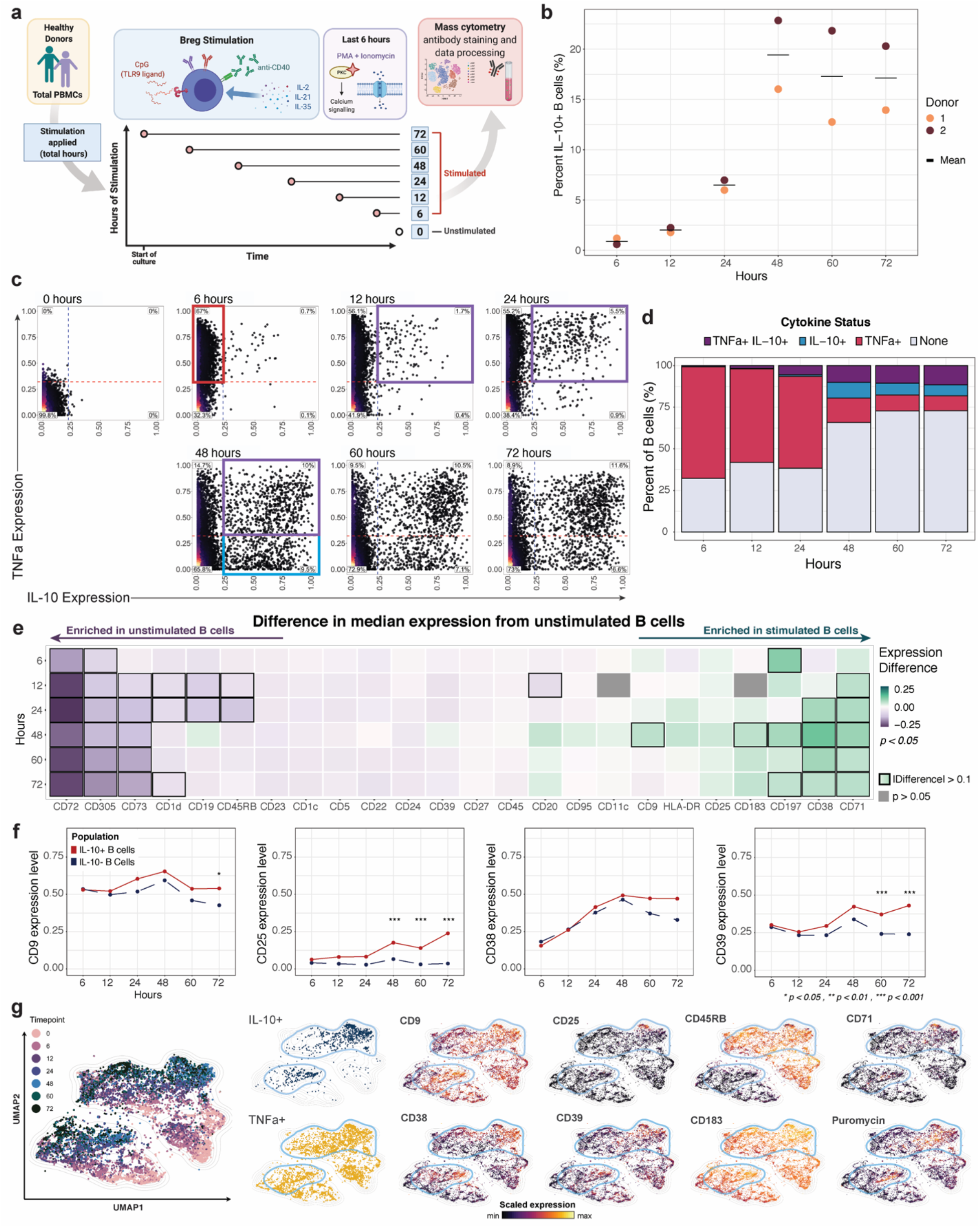
Overlapping expression of pro-inflammatory cytokines precedes IL-10 in activated B cells. **(A)** Experimental workflow **(B)** Percent of IL-10^+^ B cells by stimulation timepoint and individual (colored dots). Cross bar indicates mean percentage for each timepoint. **(C)** Biaxial plots of donor-pooled total B cells by timepoint. Red box within the plot for timepoint 6 (hours) highlights increasing TNF*a* expression. Purple boxes within the plots for timepoints 12, 24 and 48 (hours) highlight increasing proportion of B cells co-expressing TNF*a* and IL-10. Blue box within plot for timepoint 48 (hours) highlights increased proportion of B cells producing IL-10 alone. Unstimulated B cell data is depicted as timepoint 0 (hours). **(D)** Percentage of stimulated donor-pooled B cells producing TNF*a* and IL-10 by stimulation timepoint (hours). **(E)** Difference in median expression of surface markers in unstimulated versus stimulated donor-pooled B cells by timepoint (rows). Boxes in heatmap indicate markers with an absolute value of difference greater than 0.1. Expression difference only shown for markers and cytokines with a p value less than 0.05 (gray fill indicates p > 0.05). P values denote result of Kolmogorov-Smirnov test with Bonferroni multiple hypothesis correcting. **(F)** Median expression level of markers with diverging patterns between donor-pooled IL-10^+^ and IL-10^-^ B cells by timepoint. P values denote result of Kolmogorov-Smirnov test with Bonferroni multiple hypothesis correcting. * p < 0.05. *** p < 0.001. **(G)** UMAP plots of donor-pooled B cells colored by stimulation timepoint (left large panel), by positive status of indicated cytokine expression (left small panels), and by expression level of indicated surface markers (right small panels). Light blue outline highlights IL-10^+^ B cell groupings. Mass cytometry data was subsampled equally by timepoint. Unstimulated B cell data is depicted as timepoint 0. Manifold was derived using expression of only phenotypic markers.

To broadly characterize the features of *ex vivo* activated B cells at the key point of maximal IL-10 expression, we compared levels of surface markers and cytokines between stimulated and unstimulated B cell states. Total stimulated B cells recovered at 48 hours indicated significant upregulation of the activation molecules CD38, CD197 and CD183, the transferrin receptor CD71 as well as CD9, a multifunctional surface tetraspanin that impacts cellular proliferation, activation, and migration (Figure 2E). In contrast, stimulated B cells exhibited significant downregulation of the inhibitory surface proteins, CD72 and CD305, followed by the immune inhibitory control molecule CD73 and the lipid-antigen presenting molecule CD1d (Figure 2E, Figure S5). Memory markers CD27 and CD45RB and the B cell lineage marker CD19 were also downregulated upon stimulation (Figure 2E, Figure S5). These results highlight the dynamics and complexity of B cell surface molecule expression following activation and the associated challenges of applying classical gating strategies to isolate and analyze subsets of activated B cells.

### Activation-induced IL-10^+^ B cell phenotypes mirror the overall B lymphocyte population

To determine if IL-10^+^ B cells have a unique and stable phenotype that persists following activation, we compared expression of 24 surface markers on IL-10^+^ B cells as compared to IL-10^-^ stimulated B cells over a 72-hour period. The transferrin receptor CD71, the immunoregulatory markers CD9 and CD25 and the activation molecules CD39 and CD197 were significantly upregulated (p<0.05) on IL-10^+^ B cells as compared to IL-10^-^ B cells at 48 hours or later after activation (Figure 2F). However, none of these markers were enriched in IL-10^+^ B cells across all timepoints. The activation molecule CD38 was enriched among IL-10^+^ B cell populations of some 72-hour stimulation conditions of our screen (Figure 1H). In the timecourse, which assessed a single stimulation condition, CD38 was upregulated on IL-10^+^ cells only at late activation timepoints and this difference was not statistically significant as compared to IL-10^-^ stimulated B cells (Figure 2F). Embedding B cells from all stimulated timepoints by UMAP highlighted that among the transiently IL-10-associated B cell markers, CD9, CD38 and CD39 were more widely expressed across the IL-10^+^ compartment than were CD25 and CD71 (Figure 2G, blue outlines). IL-10^+^ B cell populations were also enriched for higher protein synthesis levels (indicated by puromycin) but were phenotypically indistinguishable from TNFα^+^ B cells (Figure 2G). In summary, the *ex vivo* temporal analysis revealed that expression of the pro-inflammatory cytokine TNFα precedes IL-10 induction by B cells. Furthermore, the phenotype of IL-10^+^ B cell populations is dynamic and no single surface marker uniquely defined IL-10^+^ B cells as compared to IL-10^-^ cells across stimulation timepoints.

### Conventional Breg immunophenotypes capture few IL-10-producing B cells and enrich for pro-inflammatory cytokines

A variety of phenotypic profiles have been attributed to human IL-10^+^ Bregs. To reconcile our results with previous observations (Wortel and Heidt 2017), we superimposed seven previously-defined human Breg immunophenotypes onto our stimulation and timecourse datasets and determined the fraction of total CD19^+^ B cells that they represent (Figure 3A). While each of these reported phenotypes were captured in our data, four out of seven collectively represented <2% of stimulated CD19^+^ B cells as well as <2% of IL-10^+^ CD19^+^ B cells (Figure 3A, B). The remaining three phenotypes, marked by CD1d^hi^, CD25^+^ and CD39^hi^, defined approximately 3%, 14% and 12% of stimulated CD19^+^ B cells as well as 3%, 25% and 18% of IL-10^+^ B cells, respectively, in the pooled datasets (Figure 3A). Undefined or ’other’ phenotypes represented the majority (52%) of the IL-10^+^ B cell compartment (Figure 3B). Additionally, ’other’ B cells were the predominant population among IL-10^+^ B cells in 10 of the 11 B cell stimulatory conditions in the screen and in 4 of the 6 timepoints in the CpG, anti-CD40 and rhIL-2, IL-21 and IL-35-based stimulation analysis (Figure 3C). Thus, the majority of IL-10^+^ B cells were not accounted for by previously-described Breg subsets. CD25^+^ and CD39^hi^ B cells were the only subsets enriched in the IL-10^+^ B cell population, as compared to stimulated CD19^+^ B cells (Figure 3B-C). Additionally, the relative proportions of the Breg phenotypes among total IL-10^+^ B cells varied substantially depending upon the condition and duration of stimulation (Figure 3C).

**Figure 3:**
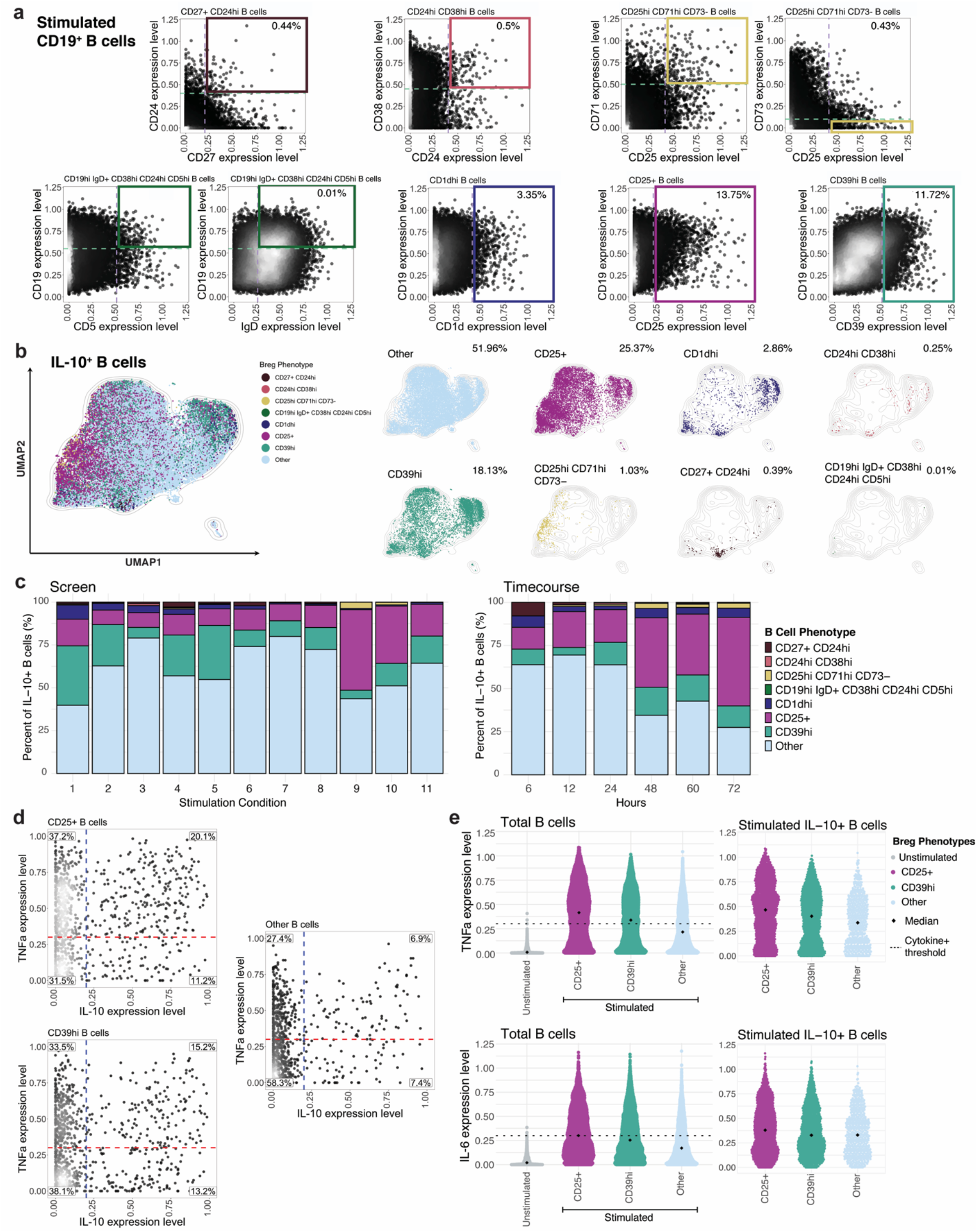
Conventional Breg immunophenotypes associated with pro-inflammatory cytokines and a limited overlap with IL-10^+^ cells. **(A)** Representative biaxial plots demonstrating gating of previously defined Breg cell surface phenotypes from the pooled stimulation condition screen and timecourse dataset. Dashed lines and solid boxes indicate thresholds of surface marker expression level within gating of CD19^+^ B cell subsets. Gating strategy for Breg phenotypes defined by >2 surface markers are indicated with 2 biaxial plots. Percent of Breg subset out of total donor-pooled CD19^+^ B cells is indicated in the upper right panel of each corresponding plot. **(B)** UMAP projection of donor-pooled IL-10^+^ B cells colored by phenotype (left panel) and with overlay of unique B cell phenotypes (right panels). Frequency of B cell phenotype among donor-pooled IL-10^+^ B cells is indicated in the upper right corner of each corresponding plot. ‘Other’ phenotype represents all remaining B cells not classified with specific surface marker-based phenotypes. Mass cytometry data was subsampled equally by condition and timepoint. Manifold was derived using expression of only phenotypic markers. **(C)** Percentage of B cell phenotypes among total donor-pooled IL-10^+^ B cells by stimulatory condition (duration of 72 hours only) from stimulation screen (left panel) and by timepoint (condition 10 only; in hours) from stimulation timecourse (right panel). **(D)** Biaxial plots of TNF*a* and IL-10 expression for CD25^+^, CD39hi and other CD19^+^ B cells from stimulation screen data. Dashed lines indicating threshold for TNF*a*^+^ status (red) and IL-10^+^ status (blue) based on unstimulated B cell expression. Percentage of B cells in each quadrant are indicated. **(E)** TNF*a* (upper panels) and IL-6 (lower panels) expression levels for total unstimulated B cells versus stimulated CD25^+^, CD39hi and other phenotype B cells (left panels) and for stimulated IL-10^+^ CD25^+^, IL-10^+^ CD39hi and IL-10^+^ other phenotype B cells (right panels). Diamond indicates median; dashed line indicates threshold for cytokine positivity based on unstimulated B cell expression.

As CD25^+^ and CD39^hi^ B cells were enriched within some IL-10-producing B cell populations, we further investigated their pro-inflammatory and immunoregulatory cytokine balance. We pooled B cell data from the stimulation screen and determined the co-expression of IL-10 and TNFα. Among CD25^+^ and CD39^hi^ B cell populations, TNFα^+^IL-10^-^ cells were highly abundant (37% and 34%, respectively), followed by TNFα^+^IL-10^+^ cells (20% and 15%, respectively) (Figure 3D). IL-10^+^TNFα^-^ cells comprised 11% and 13% of the total CD25^+^ and CD39^hi^ population, respectively. The high proportion of TNFα^+^ cells among these phenotypes, indicates that they are strongly activated B cells with pro-inflammatory features in addition to immunoregulatory cytokine secretion (Fig S6). In comparison, ’other’ B cells had a lower proportion of TNFα^+^ IL-10^-^ cells (27%) and TNFα^+^ IL-10^+^ cells (7%), indicating a less pro-inflammatory bias (Figure 3D). Both the total CD25^+^ and CD39^hi^ populations, as well as their IL-10^+^ fractions, expressed higher median levels of TNFα and IL-6 than the undefined or ’other’ B cell population (Figure 3E). Jointly, these data suggest that the IL-10-expressing CD25^+^ and CD39^hi^ B cells have a comparatively high pro-inflammatory profile relative to other IL-10^+^ B cell subsets.

Additionally, the expression of several surface molecules that have been reported as IL-10-associated in murine and human B cells (CD1c, CD1d and CD5), decreased upon B cell-specific stimulation and were not correlated with peripheral B cell IL-10 expression, regardless of stimulatory condition or duration (Figure S7-9). Proposed Breg marker CD365/TIM-1 was not detected on total unstimulated, stimulated or IL-10^+^ B cells, although CD365/TIM-1 expression was detected on *in vitro* activated CD4^+^ T cells (supp. methods, Figure S10). Collectively, previously observed Breg phenotypes comprise a limited proportion of IL-10^+^ B cell populations from healthy individuals across *ex vivo* activating conditions and timepoints and are concomitantly enriched for pro-inflammatory cytokines.

### Multiple B cell subsets can produce IL-10 and distinct polyfunctional profiles arise from each subset

While stimulation of B cells is required for robust production of IL-10, the immunophenotype of these cells is severely perturbed and not reflective of the resting state (*e.g.,* the canonical memory marker CD27 is downregulated after stimulation). To link IL-10 production to the human B cell resting state, we traced the origins of cytokine-producing cells from canonical populations using an atlas of human B cell identity established for peripheral blood and lymphoid organs (Glass et al. 2020). To understand the origins of IL-10-producing B cells, we FACS sorted six canonical B cell populations from PBMCs: CD38^+^CD24^+^ transitional, CD45RB^-^CD27^-^CD38^-^CD24^-^ naïve, CD45RB^+^CD27^-^ memory, CD45RB^+^CD27^+^ memory, CD95^+^ memory, and CD19^hi^CD11c^+^ effector cells. The purified populations were labeled with CFSE, co-cultured with a 10:1 mixture of autologous PBMCs, stimulated for 48 hours and analyzed by mass cytometry (Figure 4A). CFSE^+^ cells were then selected *in silico* to associate the resulting stimulated B cells to their original phenotypes (Good et al. 2019).

**Figure 4:**
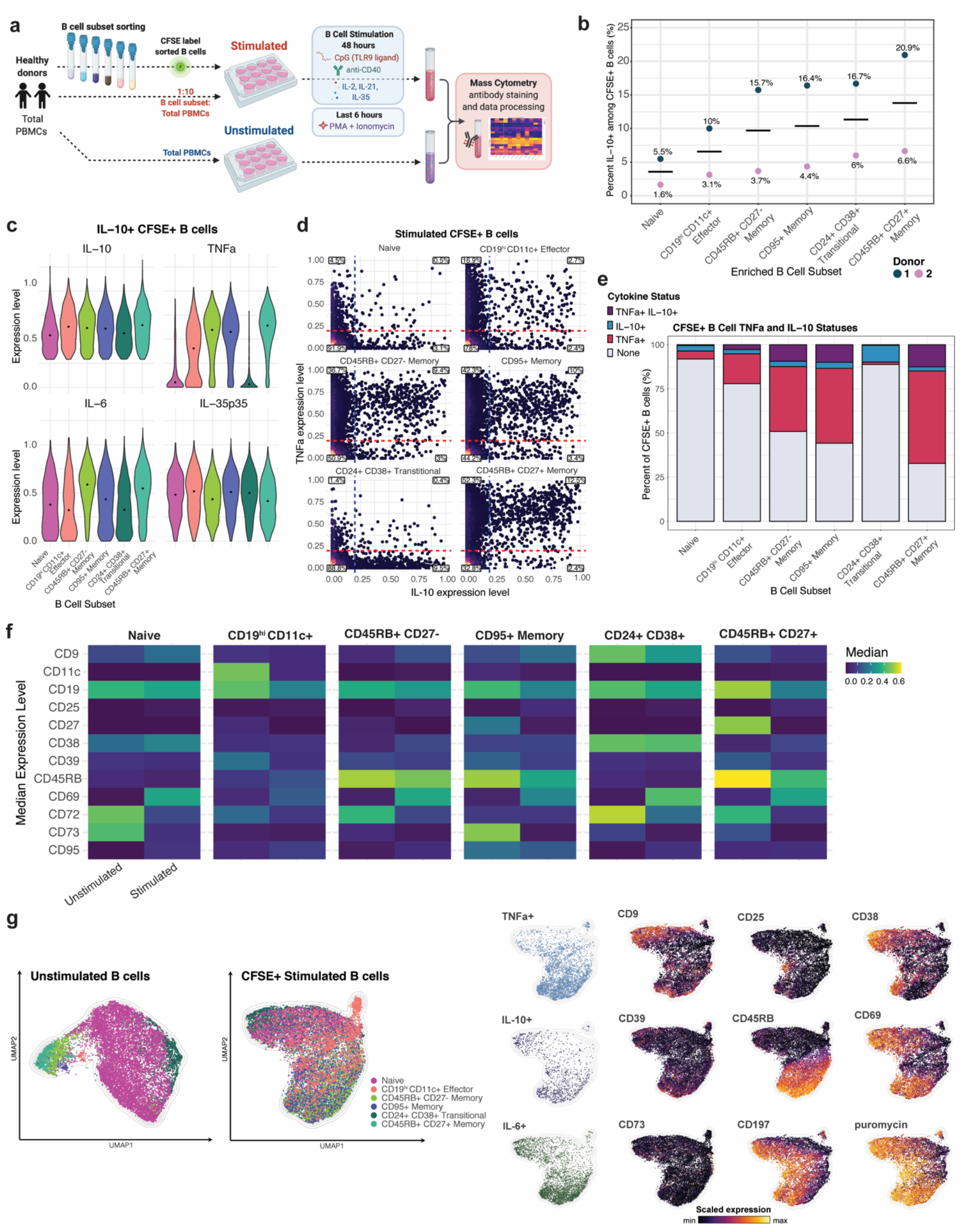
Live cell tracing reveals emergence of IL-10-producing B cells from multiple subsets and stimulation-induced phenotypic changes. **(A)** Experimental workflow **(B)** Percent IL-10^+^ cells within CFSE^+^ B cells according to resting B cell phenotype and individual (colored dots). Cross bar indicates mean percentage for each subset. **(C)** Expression levels of indicated cytokines by each of CFSE^+^ IL-10^+^ donor-pooled B cell subsets. Diamond indicates median expression. **(D)** Biaxial plots of TNF*a* and IL-10 (biaxial plots) for stimulated CFSE^+^ donor-pooled B cell subsets. Dashed lines indicating threshold for TNF*a*^+^ status (red) and IL-10^+^ status (blue) based on unstimulated B cell expression. Percentage of B cells in each quadrant is indicated. **(E)** Percentage of B cells by TNF*a* and IL-10 expression for stimulated CFSE^+^ donor-pooled B cell subsets. **(F)** Median expression level of surface markers with differential expression in unstimulated versus stimulated B cells in each of the six donor-pooled B cell subsets. **(G)** Separately-generated UMAP manifolds of unstimulated B cells (left panel) and stimulated CFSE^+^ B cells (right panels) colored by B cell subset (large left panels), by positive status of indicated cytokine expression (left small panels), and by expression level of indicated surface markers (right small panels). Equal subsampling of unstimulated and stimulated B cells from mass cytometry data. Each manifold contains 18,000 cells and was derived using expression of only phenotypic markers.

The proportion of IL-10^+^ cells that emerged from specific B cell subsets varied by individual, but the rank order of IL-10 positivity was consistent (Figure 4B). CD45RB^+^CD27^+^ memory and CD24^+^CD38^+^ transitional B cell subsets yielded the two highest proportions of IL-10^+^ B cells, relative to the naïve, CD19^hi^CD11c^+^ effector, CD45RB^+^CD27^-^ memory and CD95^+^ memory subsets (Figure 4B). Naïve B cells yielded the lowest proportion of IL-10^+^ cells (Figure 4B). This data provides the first evidence that B cells within the CD19^hi^CD11c^+^ effector memory subset produce IL-10 cytokine at the protein level (Figure 4B).

To investigate how polyfunctionality varies by subset, we compared the expression levels of cytokines produced by IL-10^+^ B cells that originated from each of the sorted B cell subsets. The median IL-10 expression level of IL-10^+^ cells within each B cell maturation stage were similar, with slightly higher expression in CD45RB^+^CD27^+^ memory cells (Figure 4C). Whereas memory B cell subsets had increased TNFa expression levels, in comparison the expression levels by IL-10^+^ cells from the naïve and transitional subsets were markedly lower. IL-10^+^ CD19^hi^CD11c^+^ effector cells also had a moderately reduced TNFα expression level (Figure 4C). The median IL-6 expression levels in IL-10^+^ cells were also higher in the memory cell subsets as compared to the effector, naïve and transitional subsets while, IL-35 p35 subunit expression levels were comparable amongst all IL-10^+^ B cell subsets (Figure 4C).

To gain further insight into the cytokine features of these B cell subsets, we specifically quantified the proportion of IL-10^-^ and TNFα-producing cells within each population. Activated CD24^+^CD38^+^ transitional B cells yielded minimal TNFα^+^ IL-10^-^ cells (<0.5%) but gave rise to the highest proportion of IL-10^+^ TNFα^-^ cells (∼10%) amongst all B cell subsets (Figure 4D-E, S11). This aligns with prior reports indicating transitional B cells had greater regulatory effects on conventional CD4^+^ T cells than naïve or memory B cells (Nova-Lamperti et al. 2016; Shabir et al. 2015; Newell et al. 2010; Cherukuri et al. 2014, 2017, 2021; Hasan et al. 2019; Laguna-Goya et al. 2020). Activation of naïve B cells yielded relatively few TNFα^+^ or IL-10^+^ cells, such that most cells did not produce either cytokine (∼92%) (Figure 4D-E, S11). This suggests naïve B cell have a higher activation threshold and/or lower responsiveness to Breg-specific stimulatory conditions, mirroring our recent findings that naïve cells are more anergic than other B cell subsets (Glass et al. 2020). CD45RB^+^CD27^+^ memory B cells had the highest proportion of TNFα^+^ IL-10^-^ cells (∼52%) and TNFα^+^ IL-10^+^ cells (∼13%), and the lowest proportion of TNFα^-^ IL-10^-^ cells (∼33%) (Figure 4D-E, S11). Overall, memory B cells, particularly the CD45RB^+^CD27^+^ B cell subset, are more skewed towards pro-inflammatory responses than other B cell maturation states under the same *in vitro* activation conditions.

To further characterize the phenotype of the IL-10-producing cells that emerged from each B cells subset, we assessed the expression of cell surface proteins associated with cellular activation and IL-10 production. As expected, the activation markers, CD69, and to a lesser extent CD25, were upregulated by all B cell subsets confirming their responsiveness to the stimuli utilized (Figure 4F). CD9, which has been associated with IL-10^+^ Bregs, was upregulated within the naïve, CD24^+^ CD38^+^ transitional, CD45RB^+^ CD27^-^ memory and CD95^+^ memory subsets. Median expression of the activation molecule CD38 was higher for the naïve, CD45RB^+^ CD27^-^ memory and CD45RB^+^ CD27^+^ memory subsets. Interestingly, CD39 was downregulated, but still detectable, on all B cell subsets. The reduction in CD39 expression was surprising given previously-reported correlations between IL-10 and CD39 expression (Hasan et al. 2019, 2021). The canonical memory marker CD27 was downregulated by all memory B cell subsets, while the median expression levels of CD45RB decreased only slightly within 3 of the 4 memory B cell subsets. CD72 and CD73 were markedly downregulated by all B cell subsets, resulting in complete loss of expression in the case of CD73 (Figure 4F).

Importantly, the requirement of Breg-specific stimulation for the detection of IL-10 producing B cells led to substantial phenotypic changes within all B cell subsets analyzed (Figure 4F). Upon stimulation, surface marker expression changes within distinct B cell subsets caused these populations to become phenotypically similar and largely indistinguishable, thereby precluding the isolation of the subsets by resting B cell phenotype gating strategies. UMAP visualizations of unstimulated and stimulated B cell subsets reveal the phenotypic changes and loss of distinct surface features (*i.e.,* cells colored by resting B cell subset become highly overlapping within plot) amongst the subsets upon stimulation (Figure 4G). Furthermore, stimulated IL-10^+^ B cells within all subsets showed moderate positive correlations with expression of the surface markers CD9, CD39, CD38, CD69, CD197 as well as the pro-inflammatory cytokines and protein synthesis activity (Figure 4G, S12). Together, tracking of B cell subsets through *ex vivo* stimulation showed that each B cell subset can potentially produce IL-10, yet each subset’s IL-10^+^ population bears unique, concurrent proinflammatory cytokine profiles.

### IL-10-producing B cells potential is increased in operationally tolerant liver allograft recipients

To assess the IL-10 producing B cells in transplantation, we stimulated the PBMCs of liver transplant recipients receiving maintenance IS, operationally allograft tolerant liver transplant recipients off IS for >1 year, and healthy controls and assessed the immunosuppressive IL-10 versus proinflammatory cytokine production by their B cell compartments (supplementary table 1, Figure 5A). We determined that operationally graft tolerant liver transplant recipients who had maintained their allograft in the absence of IS had a significantly higher proportion of B cells capable of IL-10 production (p=0.02) and overall elevated expression levels of IL-10 as compared to liver transplant recipient controls (Figure 5B, S13, supplementary table 2). In contrast, there was no significant difference in the proportion of TNFα^+^ or IL-6^+^ B cells between the groups (Figure 5B, S14). Operationally graft tolerant patients and healthy control subjects had higher proportions of IL-10^+^ TNFα^+^ B cells (16% and 18%, respectively) and IL-10^+^ TNFα^-^ B cells (8% and 10%, respectively), as compared to control transplant recipients (6% IL-10^+^ TNFα^+^ and 2% IL-10^+^ TNFα^-^) (Figure 5C). The proportions of IL-10^-^ TNFα^+^ B cell were similar between the two liver transplant recipient groups, and lower in the healthy control group. We quantified the log2 ratio of IL-10^+^:TNFα^+^ total B cells and found the operationally tolerant cohort had an approximately 2-fold higher ratio than the transplant control cohort, suggesting a greater immunoregulatory bias of their B cell compartment (Figure 5D).

**Figure 5:**
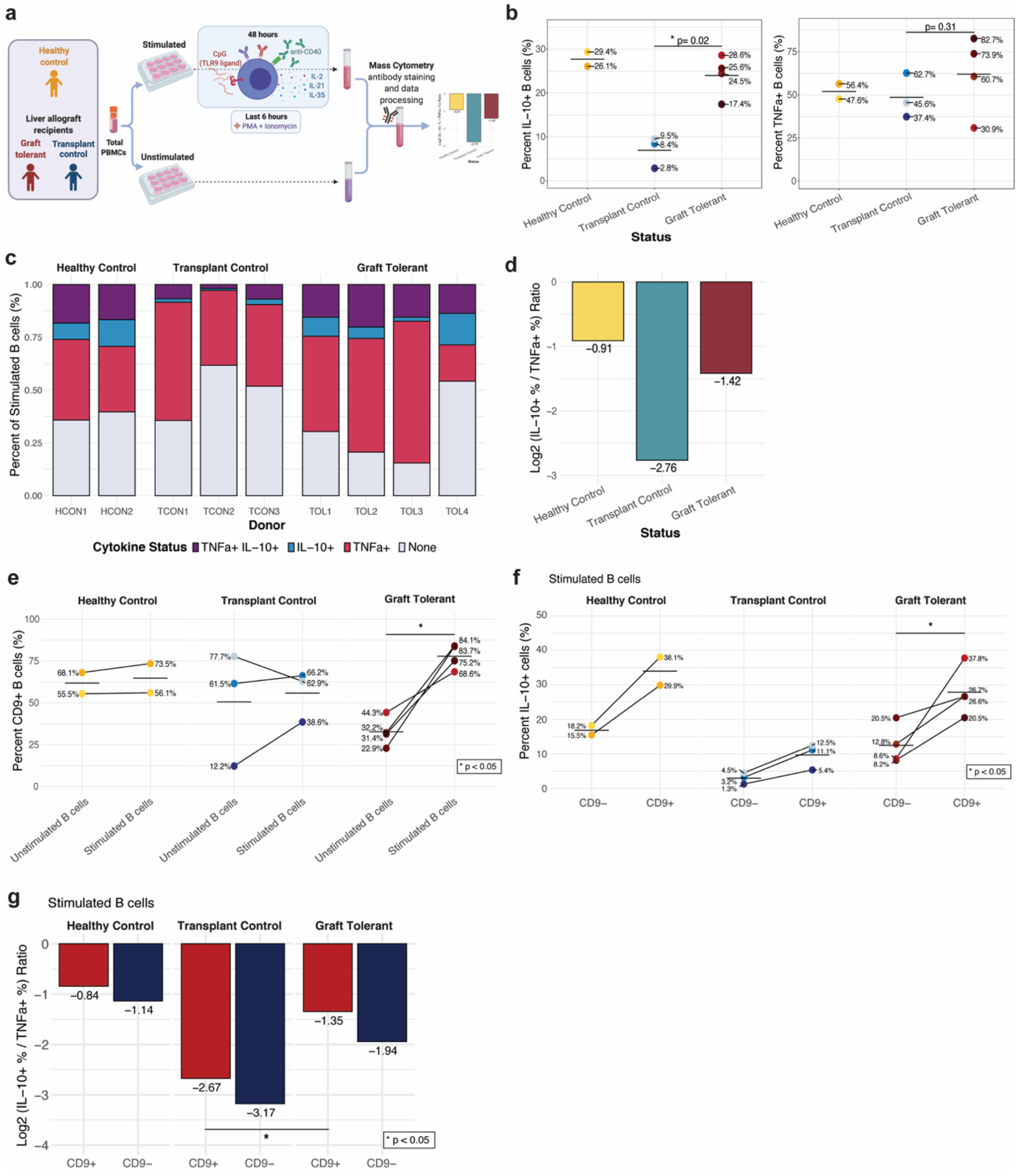
Operationally graft tolerant liver transplant recipients are enriched for IL-10-producing B cells. **(A)** Experimental workflow to analyze B cells of liver transplant recipients on immunosuppression (IS), operationally allograft tolerant liver transplant recipients off IS for >1 year, and healthy controls (no transplant). **(B)** Percent IL-10^+^ (left plot) and TNF*a*^+^ (right plot) B cells by group and individual (colored dots). Cross bar indicates mean percentage for each group. P values denote result of Wilcoxon rank sum test. * p < 0.05. **(C)** Percentage of B cells with each TNF*a* and/or IL-10 expression status among total stimulated B cells by group (panels) and individual (columns). **(D)** Log2 ratio of percentage IL-10^+^ to TNF*a*^+^ donor-pooled B cells by group. **(E)** Percentage of CD9^+^ B cells among unstimulated and stimulated B cells by group (panel) and individual (colored dots). P value denotes result of Wilcoxon signed-rank test. * p < 0.05 **(F)** Percentage of IL-10^+^ cells among CD9^-^ and CD9^+^ stimulated B cells by group (panel) and individual (colored dots). P value denotes result of Wilcoxon signed-rank test. * p < 0.05 **(G)** Log2 ratio of percentage IL-10^+^ to TNF*a*^+^ cells for CD9^+^ (red) and CD9^-^ (blue) donor-pooled B cells by group. * p < 0.05

To determine whether there were unique features that define the IL-10-producing B cells in clinical transplant recipients, we examined cell phenotypes within patient groups. Notably, CD9^+^ cells were uniquely and significantly (p<0.05) increased within stimulated B cells of operationally tolerant liver allograft recipients (Figure 5E, S15). The CD9+ B cell populations of operationally tolerant graft recipients had a significantly higher proportion of IL-10^+^ cells as compared to CD9^-^ populations (p<0.05). There was a trend towards an increased proportion of IL-10^+^ cells in CD9^+^ B cell populations in healthy controls and transplant controls, but the differences were not significant (Figure 5F). CD9^+^ B cells also had a higher log2 ratio of IL-10^+^:TNFα^+^ B cells as compared to CD9^-^ populations within all groups (Figure 5G). The stimulated CD9^+^ B cell population of operationally tolerant transplant recipients had a significantly higher ratio of IL-10^+^:TNFα^+^ cells (*i.e.,* was enriched for IL-10 production) as compared to transplant controls (p<0.05). However, CD9 expression was not detected on all IL-10^+^ B cells in any of the subjects, nor did all CD9^+^ B cells produce IL-10 (Figure 5F, S16). In summary, operationally graft tolerant liver transplant recipients exhibited a higher frequency and magnitude of IL-10-production in B cells with a corresponding increase in the trafficking molecule CD9. Together this suggests that, despite heterogenous immunophenotypes, the IL-10-producing state of circulating B cells in a context of reduced proinflammatory cytokine secretion may contribute to immune tolerance and allograft homeostasis.

## Discussion

Here, we employed high-dimensional, deep immune profiling to show that human peripheral IL-10-producing B cells exhibit highly activated, yet heterogenous B cell phenotypes. Through comprehensive assessment across an array of *ex vivo* activating conditions using TLR ligands, anti-CD40 and cytokines, and timepoints, as well as through precise CFSE-based B cell subset tracking, we demonstrate that IL-10^+^ B cell surface and cytokine profiles are diverse and strongly influenced by the method and duration of activation. Furthermore, we show that a variety of resting, canonical B cell subsets can give rise to IL-10^+^ B cells. In fact, no single, unifying phenotype captured the majority of IL-10^+^ B cells. Instead, we found that IL-10^+^ B cells extend well beyond those formerly proposed, discrete Breg phenotype cells, such as CD39^hi^, CD25^+^ and CD1d^hi^ B cells (summarized by *Wortel and Heidt 2017)* (Wortel and Heidt 2017). Moreover, seven previously described Breg subsets represented only a minor proportion of the total IL-10-producing B cell compartment in peripheral blood of humans and produced high levels of proinflammatory cytokines. Notably, the range of B cell-specific stimulatory conditions applied in our experiments included *ex vivo* conditions utilized in multiple prior Breg studies. These findings collectively demonstrate that IL-10^+^ B cells or Bregs lack the features of a dedicated immunological subset based on current definitions and represent a dynamic, impermanent, and regulatory-skewed state enriched within the transitional B cell subset. These findings confirm and extend the work of *Lighaam et al.,* who showed that *ex vivo* IL-10 production is the transient property of an activated B cell state (Lighaam et al. 2018).

Based on our data, we hypothesize that the regulatory capacity of B cells may be more accurately quantified by their cytokine co-expression profiles than solely by the detection of IL-10 expression. A significant portion of IL-10^+^ B cells co-expressed IL-6 and TNFα, suggesting that IL-10^+^ B cells are not a distinct, monofunctional population. Prior studies established that human IL-10^+^ peripheral B cells co-express the proinflammatory cytokines TNFα and IL-6 (Lighaam et al. 2018; Cherukuri et al. 2014), and that transitional B cells have a higher ratio of IL-10:TNFα. These observations coincided with the finding that transitional B cells demonstrate significantly enhanced immunoregulatory activity on CD4^+^ T cells as compared to other stimulated B cell subsets, suggesting that IL-10:TNFα ratios reflect a meaningful metric of immunoregulatory capacity (Nova-Lamperti et al. 2016; Shabir et al. 2015; Newell et al. 2010; Cherukuri et al. 2014, 2017, 2021; Hasan et al. 2019; Laguna-Goya et al. 2020). In the present study, we observed that transitional B cells were the only subset with a positive IL-10:TNFα log ratio amongst B cell subsets, suggesting that the immunoregulatory potential of B cells is diminished upon maturation and differentiation. Further, the immunoregulatory bias of transitional B cells may result from their recent emigration from the bone marrow, in which inflammation during B cell lymphopoiesis would be undesirable.

Our ability to understand the heterogeneity of the IL-10^+^ B cell compartment hinged on comprehensively assessing *ex vivo* stimulation conditions and durations, while excluding those previously tested modes of stimulation, such as BCR crosslinking, with contradictory reports on *ex vivo* induction of IL-10 from B cells (Iwata et al. 2011; Lighaam et al. 2018; Bankó et al. 2017). Nevertheless, several unexplored conditions, such as stimulation with R848, anti-CD40 and exogenous IL-4 and IL-10, were tested to evaluate B cell cytokine production and immunophenotypes. Our results indicate that CpG stimulation combined with CD40 activation and addition of exogenous human IL-2, IL-21 and IL-35 of human PBMCs was the strongest inducer of IL-10 expression in B cells and the peak of this IL-10 production was at 48 hours of *ex vivo* stimulation. In a related study, TLR9-based activation with CpG yielded the highest proportion of IL-10^+^ cells and highest levels of IL-10 expression in adult B cells, as compared to all other TLR agonists, alongside CD40 activation (Iwata et al. 2011). Although functional impacts of B cell subsets on T cells were not investigated in the current study, prior studies have established that stimulation of human B cells with CpG TLR9 agonist and CD40 activation, in addition to exogenous cytokines in some studies, induces IL-10 production and immunosuppressive function from specific IL-10^+^ B cell subsets (Iwata et al. 2011; Cherukuri et al. 2014, 2021). These immunosuppressive activities include inhibition of proliferation, activation and proinflammatory cytokine (IFNg, TNFα) expression from conventional CD4^+^ T cell in humans (Iwata et al. 2011; Cherukuri et al. 2014, 2021; Bouaziz et al. 2010). The overlap of the focal activating cocktail used in the later part of our study and in these prior works, reinforces the notion that IL-10^+^ B cell populations profiled here likely possess immunoregulatory activity.

Similar to other studies in humans, we could not compare IL-10 production by B cells from multiple tissues in the same individual. Tissue-resident IL-10-producing B cells may share characteristics with those circulating in peripheral blood. Along these lines, a prior study established that B cells derived from newborn cord blood and adult tonsil and spleen tissues exhibited the highest IL-10 production after stimulation with CpG and CD40 activation for 48 hours in a screen of *ex vivo* stimulations (Iwata et al. 2011). This previous work and the data from the current study further highlight the necessity and potency of TLR9 agonist and CD40 activation in induction of IL-10 from human B cells as well as the low or undetectable levels of IL-10 from *in vivo* sampled cells. A comprehensive and high-dimensional assessment of unique, tissue-defined IL-10^+^ B cell phenotypes would be a useful resource and follow-up to the present study.

The basis of our interrogation of human IL-10^+^ B cells utilized PBMCs from healthy control individuals, to achieve an analysis that was not biased by immune dysregulation and, instead, focused on immune homeostasis. To extend this to a clinical scenario, we analyzed peripheral IL-10^+^ B cells in operationally tolerant liver transplant recipients to investigate whether these cells could be contributing to the establishment or maintenance of allograft tolerance. Most studies of clinical transplantation have focused on the levels of Bregs as indicators of impending rejection or long-term outcome, rather than on transplant tolerance. The operationally tolerant subjects in our study had been off IS for over one year and had not experienced a rejection episode during that period. We analyzed operationally tolerant liver allograft recipients, as it has been suggested that liver transplant recipients include a larger proportion, compared to other solid organ recipients, of individuals who are spontaneously tolerant and could maintain their allograft in the absence of IS (Feng 2016; Lerut and Sanchez-Fueyo 2006; Dai et al. 2020). We found that operationally allograft tolerant liver transplant recipients have a higher proportion of IL-10-producing B cells and elevated expression levels of IL-10 by B cells (similar to healthy control subjects) as compared to transplant recipient controls. It should be noted that the transplant controls subjects were receiving IS, while graft-tolerant subjects were not, which could contribute to the differences observed. Yet, the same trend in B cell functional differences seen with IL-10 was not observed with the pro-inflammatory cytokines, TNFα and IL-6. We additionally found that stimulated, total CD19^+^ B cells of operationally tolerant liver recipients had a markedly higher IL-10:TNFα ratio than transplant controls and CD9 expression showed a unique and significant correlation with IL-10 production by peripheral B cells from tolerant subjects. Thus, the B cell compartment of the operationally tolerant liver transplant recipient cohort is collectively skewed towards an immunoregulatory state, which is associated with, but not defined by, higher CD9 expression. This raises the possibility that B cells are contributing to the natural allograft tolerance observed in this cohort. Importantly, this study provides an analysis of IL-10-producing B cells in a clinical cohort that is defined by a unique state of immune tolerance, and diverges from most clinical studies, which described B cells in a disease state defined by immune dysregulation.

The tetraspanin CD9 is well-established as a marker of murine Bregs with immunosuppressive activity and IL-10 production (Brosseau et al. 2018; Matsushita et al. 2016; Braza et al. 2015; Sun et al. 2015). In humans, B cells of healthy individuals are known to co-express CD9 and the immunoregulatory cytokine IL-10 upon activation, and CD9 expression is enriched in CD24^hi^CD38^hi^ transitional B cells (Hasan et al. 2019). Here, we report for the first time a significant correlation between expression of CD9 and IL-10 by stimulated B cells of operationally liver allograft tolerant transplant recipients and a higher IL-10:TNFα ratio in CD9^+^ cells versus CD9^-^ cells (**Figure 5**). However, not all CD9^+^ B cells produced IL-10 and, in fact, 38-85% of IL-10^-^ stimulated B cells were CD9^+^. Interestingly, the positive correlation between IL-10 and CD9 expression by B cells was not significant within healthy individuals until 72 hours after *ex vivo* stimulation. We further detected the highest level of CD9 expression in transitional B cells as compared to other subsets, when examining unstimulated, stimulated and IL-10^+^ B cell states in healthy individuals. This suggest that CD9 is associated with the B cell IL-10 production, but neither a requirement nor an instrumental or fixed surface marker of IL-10^+^ B cells.

In conclusion, we have utilized high-dimensional cytometry to reveal the phenotypic diversity and origins of human IL-10^+^ B cells and demonstrate the absence of a single, canonical subset of IL-10^+^ B cells in the peripheral blood of healthy humans and liver transplant recipients. Our findings provide a comprehensive overview of the phenotype and cytokine expression profiles of IL-10^+^ human B cells across a range of *ex vivo* activating conditions and durations. This study also emphasizes the importance of assessing B cell polyfunctionality alongside IL-10 expression to accurately distinguish between proinflammatory and immunoregulatory-skewed populations. The dynamic and heterogenous nature of IL-10-producing B cells, and the enrichment of the IL-10^+^ cell state among transitional B cells should be considered when analyzing total or sorted IL-10^+^ B cell populations. Despite the variable origins and potential impermanence of IL-10^+^ B cells, the impact of the IL-10^+^ activated B cell state has been clearly associated with immunoregulatory activity and outcomes in human disease and allograft responses. This work should serve as a resource to future studies on IL-10-producing B cells and their immunoregulatory contributions in humans.

## Methods

### Study design

The purpose of this study was to provide comprehensive and high-dimensional analyses of the phenotypic heterogeneity and polyfunctionality of IL-10^+^ B cells in the context of healthy individuals and operationally graft tolerant liver transplant recipients. To this end, we tested *in vitro* stimulation conditions and durations of human PBMC samples for optimal B cell IL-10 expression. We assessed the origins of IL-10-producing cells by sorting six defined B cell subsets and tracked these cells through the Breg-inducing activation process with CFSE labeling. Finally, we applied optimum *in vitro* conditions identified from prior experiments to PBMCs from immunosuppressed organ transplant recipients and operationally graft tolerant transplant recipients to investigate the IL-10^+^ B cell profile in these patient groups.

### Human ethics and blood collection

Deidentified human blood was obtained from healthy adult donors (Stanford Blood Center, n=8). Blood samples were also collected from pediatric and adult liver transplant recipients (n=7) at Lucile Packard Children’s Hospital and Stanford Hospital. All samples were obtained under informed consent and in accordance with Stanford’s Institutional Review Board. PBMCs (lymphocytes) were purified from fresh whole blood samples using Ficoll-Paque (GE Healthcare) gradients. Healthy blood donor and transplant recipient patient cells were processed immediately for *in vitro* stimulations or cryopreserved in pure fetal bovine serum with 10% dimethyl sulfoxide at −80°C overnight and −196°C long-term. *Cell culture and stimulation of peripheral blood mononuclear cells*

Total PBMCs were suspended in RPMI 1640 supplemented with 10% fetal bovine serum, 1% penicillin-streptomycin, L-glutamine, HEPES, sodium pyruvate, glucose and 2-Mercaptoethanol (Gibco; 50µM) at 1x10^6^ cells/mL and rested for 30 minutes. In the stimulation screen experiment, rested PBMCs were stimulated for 72 hours with TLR ligand, with or without CD40 activation and with or without a variety of exogenous recombinant human cytokines, as outlined in Figure 1A and 1B. Stimulation reagents and concentrations are outlined in supplementary table 3. In all stimulations, phorbol 12-myristate 13-acetate (PMA; 50 ng/mL), ionomycin (1 µg/mL), and Brefeldin A (BFA;1x) were added for the final 5 hours of culture. In the stimulation timecourse experiment, rested PBMCs were stimulated with CpG ODN2006 (10 µg/mL) and anti-CD40 (500 ng/mL) as well as exogenous recombinant human IL-2 (600 IU/mL), IL-21 (100 ng/mL) and IL-35 (20 ng/mL) for 6, 12, 24, 48, 60 and 72 hours (outlined in Figure 2A), with addition of PMA, ionomycin, and BFA in the final 5 hours. In the B cell sorting and transplant recipient cohort experiments, the same Breg-specific stimulation with CpG, anti-CD40 and exogenous cytokines as well as PMA/ionomycin and BFA was applied for a total of 48 hours (outlined in Figure 4A, 5A).

### Mass cytometry processing and acquisition

Antibody conjugation, staining, and data acquisition were performed as previously described (Hartmann et al. 2018). Briefly, metal-isotope labeled antibodies used in this study were conjugated using the MaxPar X8 Antibody Labeling kit per manufacturer instruction (Fluidigm) or were purchased from Fluidigm pre-conjugated. Each conjugated antibody was quality checked and titrated to optimal staining concentration using a combination of primary human cells (Figure S7-10). PBMCs from cultures were spiked with a tri-molecular biosynthesis cocktail, consisting of 5-Bromouridine (BRU; Sigma, 5mM), 5-Iodo-2-deoxyuridine (IdU; Sigma, 100uM) and puromycin (Sigma, 10 μg/mL), and incubated for 30 minutes (Kimmey et al. 2019), then washed in cell staining media (CSM; PBS with 0.5% BSA and 0.02% sodium azide and benzonase 25x10^8^ U/mL (Sigma)) and resuspended in 0.5µM cisplatin in phosphate-buffered saline (PBS) for 5 minutes to label non-viable cells (Sigma). Cells were washed in CSM and fixed with 1.6% PFA in PBS for 10 minutes at room temperature (RT). Fixed cells were Palladium barcoded (Fluidigm) as previously described (*94*). Barcoded cells were then suspended in TruStain FC blocker (Biolegend) for 10 minutes at RT and washed in CSM prior to staining. All surface staining was performed in CSM for 30 minutes at RT. Cells were washed in CSM and permeabilized with 0.02% saponin (Sigma) in CSM for 30 minutes on ice. Intracellular and anti-biotin staining was performed in 0.02% saponin in CSM for 1 hour at RT. Before acquisition, samples were washed in CSM and resuspended in intercalation solution (1.6% PFA in PBS, 0.02% saponin (Sigma) and 0.5µM iridium-intercalator (Fluidigm)) for 1 hour at RT or overnight at 4°C to label DNA. Before acquisition, samples were washed once in CSM and twice in ddH2O. All samples were filtered through a 35µm nylon mesh cell strainer, resuspended at 1x10^6^ cells/mL in ddH2O supplemented with 1x EQ four element calibration beads (Fluidigm), and acquired on a CyTOF2 mass cytometer (Fluidigm) using the Super Sampler injection system (Victorian Airship).

### Sorting of B cell subsets and carboxyfluorescein diacetate succinimidyl ester-based cell tracking

Total PBMCs were isolated from Trima Accel leukocyte reduction system (LRS) chambers as described above and then Fc blocked and subjected to magnetic lineage depletion according to the manufacturer’s instructions using BD Streptavidin Particles Plus and the BD IMag Cell Separation Magnet (BD Biosciences) with a cocktail of biotinylated antibodies consisting of CD3, CD7, CD15, CD33, CD56, CD61, and CD235ab (Biolegend). The cells bound by the biotinylated antibody cocktail were captured by streptavidin particles for depletion. The remaining B cell-enriched suspensions were stained for surface antigens (antibodies listed in supplementary table 4) in CSM in the dark for 30 minutes on ice and then washed in CSM. Stained cell suspensions were incubated with eFluor 780 viability stain (eBioscience; 1:1000 dilution) in the dark for 5 minutes on ice to label non-viable cells, then immediately washed in PBS and transferred for cell sorting on a BD Biosciences FACS Aria II equipped with violet, blue and red lasers (Stanford Blood Center, Flow Cytometry Core). Within the cell sorting phase, total B cells were defined by surface expression as CD20^+^ and B cell subsets were identified as CD24^-^ CD38^-^ naïve, CD24^+^CD38^+^ transitional, CD45RB^+^CD27^+^ memory, CD45RB^+^CD27^-^ memory, CD95^+^ memory, and CD11^+^ effector memory cells. 200,000 cells from each B cell subset were sorted into pure FBS on ice and immediately washed 3 times in 37°C PBS and labeled with CellTrace carboxyfluorescein succinimidyl ester (CFSE) Cell Proliferation Kit (Invitrogen; 1µM) according to the manufacturer’s instructions, followed by an RPMI wash and final resuspension in 37°C complete RPMI (as described above). The sort gating strategies for individual B cell subsets and validation of CFSE-labeling of these subsets are shown in Figure S17. Each CFSE-labeled B cell subset was added to two replicate cultures with autologous donor total PBMCs at a 1:10 final concentration (sorted B cell subsets:PBMCs) and 2x10^6^ cells/mL. A Breg-specific stimulation, as described above, was applied to final cultures for 48 hours (outlined in Figure 4A).

### Mass cytometry data pre-processing

Data processing were performed as previously described (Glass et al. 2020). Briefly, acquired samples were bead-normalized using MATLAB-based software as previously described (Finck et al. 2013) and debarcoded using MATLAB-based software (Zunder et al. 2015). Normalized data was then uploaded onto CellEngine (*cellengine.com, Primity Bio, Fremont, CA*) for singlet (Tsai et al. 2020) and B cell gating (Figure S18). Gated data was downloaded and further processed with the R programming language (http://www.r-project.org) and Bioconductor (http://www.bioconductor.org) software. Data was transformed with an inverse hyperbolic sine (asinh) transformation with a cofactor of 5. Each marker was scaled to the 99.9th percentile of expression of all cells in that experiment. Cytokine positive thresholds for B cells were defined by the 99.9th percentile of expression for each cytokine in unstimulated B cells within each experiment. Batch correction was not applied as all samples within an experiment were barcoded and acquired in the same mass cytometry run, except for individual samples in the B cell subset sorting experiment. For the B cell sorting experiment, batch effect was minimal, so markers were 99.9^th^ percentile scaled by batch and then combined for analysis.

### Data visualization

To visualize co-expression of molecules measured on different B cell populations in the stimulation screen, timecourse and B cell sorting experiments, cells labeled with the same mass cytometry panels were subsampled (equally by B cell population, for the screen analysis) and plotted on a UMAP plot using the umap package in R (Figure 1E, 2E, 3B, 4G). The plots were generated based on expression of all phenotypic markers expressed on B cells. In the stimulation screen and B cell phenotype analysis, the UMAP plots were additionally based on cytokine expression (Figure 1E, 3B). Isotype was not used to generate maps to prevent artificial separation of phenotypically similar cells. Subsampling by B cell population facilitated visualization of heterogeneity without the map being dominated by the most abundant populations.

For activated B cell state analysis, stimulated B cells were over-clustered into 100 clusters using FlowSOM with all detectable surface molecules as input. Clusters were then hierarchically clustered based on expression of B cell surface molecules and isotype.

Unstimulated B cells were over-clustered with FlowSOM and manually assigned to cell subsets, as described previously (Glass et al. 2020).

### Statistics

All statistical tests for differences in distribution of markers and cytokines between B cell populations from mass cytometry data were performed on equally subsampled populations using the KS test. This non-parametric test determines the equality of two continuous distributions. All P values were corrected by the Bonferroni method, the most conservative of multiple hypothesis correction approaches. For comparisons of single marker expression levels between B cell populations and frequencies of single B cell populations between conditions, timepoints or clinical groups, the Wilcoxon rank sum test was performed with Bonferroni correction.

## Acknowledgments

We thank the flow cytometry core staff at Stanford Blood Center for guidance and assistance with cell sorting experiments. We also thank P. Favaro, J. Pena and J. Toh for guidance with B cell sorting experiment plans, F. Hartmann, D. Mrdjen and J. Harden for guidance with mass cytometry experiments and T. Bruce, S. Greenbaum, S. Kimmey, E. McCaffery and A. Calderon for experimental and reagent contributions. Lastly, we thank B. Sahaf and K. Bruton for advice on optimization of human B cell culture conditions. Figures were created using BioRender (http://www.biorender.com) and Illustrator (Adobe).

## Author contributions

Conceptualization: MCG, DRG, OMM

Experimental Design: MCG, DRG, JPO, BM

Investigation: MCG, DRG, JPO, BM

Visualization: MCG, DRG

Funding acquisition: OMM, SCB, SMK, COE

Supervision: OMM, SCB

Writing – original draft: MCG, OMM

Writing – review & editing: DRG, COE, SMK, SCB

## Code and data availability

All data are available in the main text or the supplementary materials.

