## Supplementary Materials for "Human IL-10-producing B cells have diverse states induced from multiple B cell subsets"

**Supplementary Methods**:

*CD365/TIM-1 Mass Cytometry Antibody Validation*

$CD4^+$ T cell stimulation method consisted of total PBMCs treated with 3 days with plate-bound anti-CD3 (10 ug/mL) and anti-CD28 (5 ug/mL) and soluble human IL-2 (50 IU/mL) and IL-4 (12.5 ug/mL) and addition of Brefeldin A (1x) in the final 5 hours. Breg stimulation method consisted of total PBMCs treated with 2 days with CpG ODN2006 (10 ug/mL) and anti-CD40 (500 ng/mL) as well as exogenous recombinant human IL-2 (600 IU/mL), IL-21 (100 ng/mL) and IL-35 (20 ng/mL) with addition of PMA (50 ng/mL), ionomycin (1 ug/mL), and Brefeldin A (1x) in the final 5 hours.

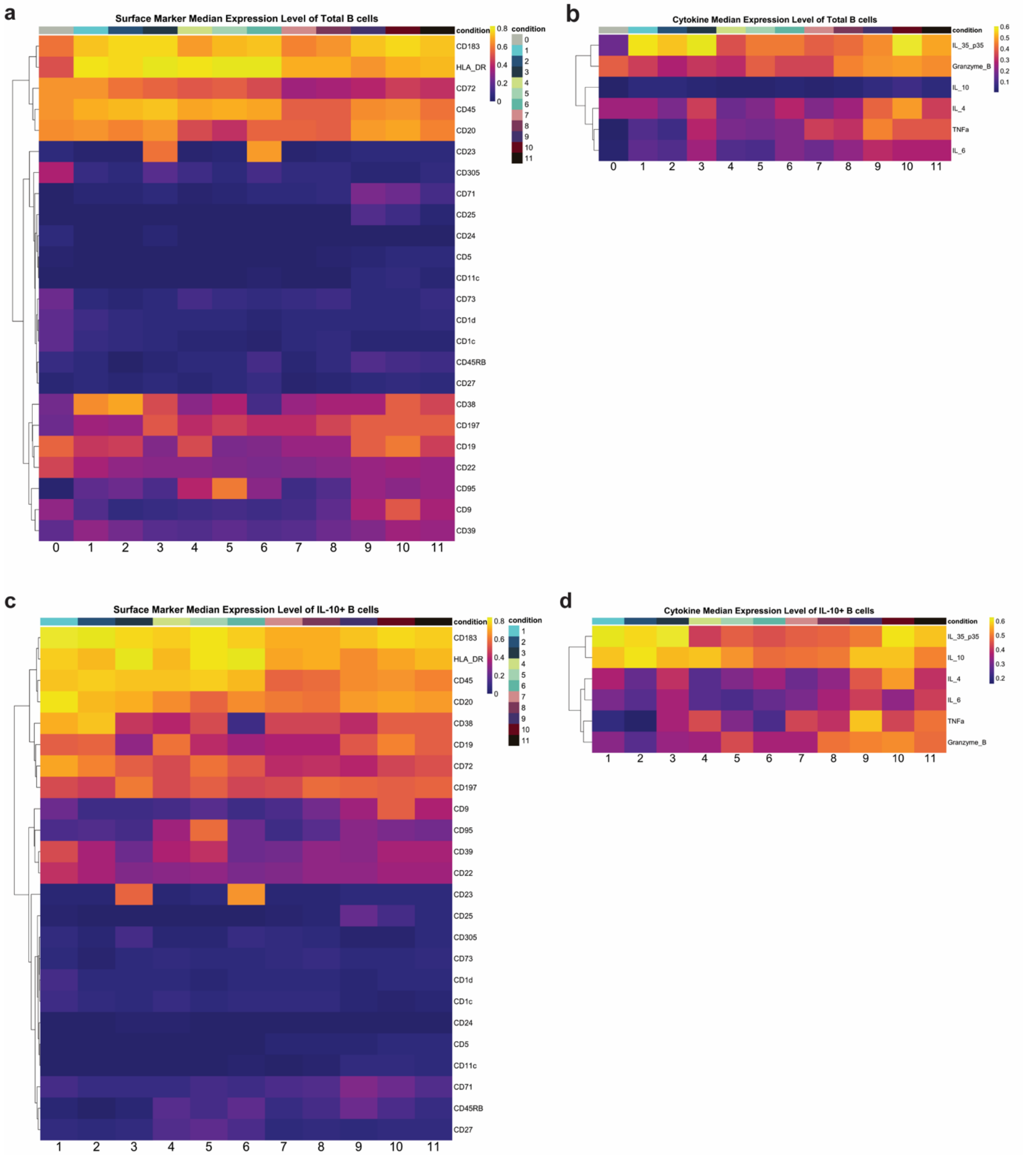


**Figure S1**. Surface marker and cytokine expression heatmaps of total and IL-10$^{+}$ B cells from stimulation screen.

**(A)** Median surface marker expression of total B cells in the unstimulated condition (0) and each stimulated condition (1-11).

**(B)** Median cytokine expression of total B cells in the unstimulated condition (0) and each stimulated condition (1-11).

**(C)** Median surface marker expression of IL-$10^{+}$ B cells from stimulatory conditions 1-11.

**(D)** Median cytokine expression of IL-$10^{+}$ B cells from stimulatory conditions 1-11.

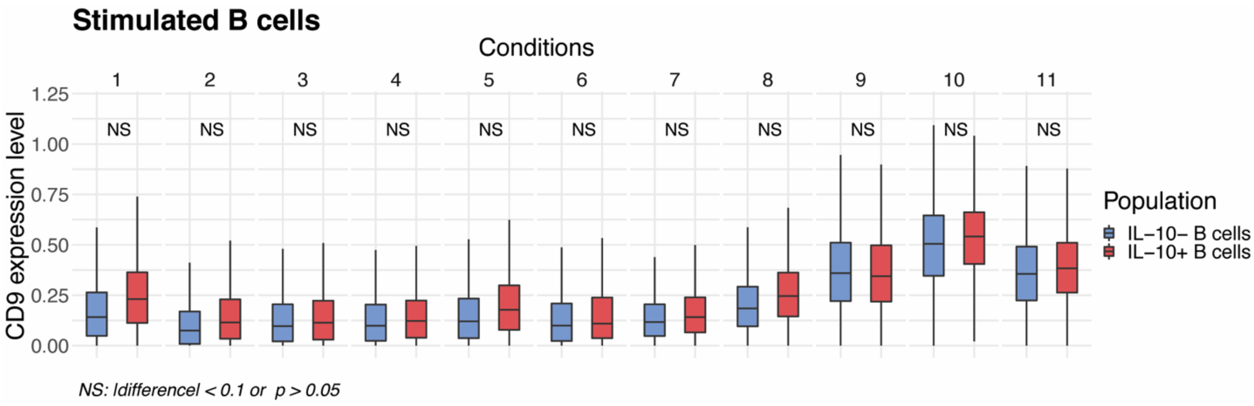


**Figure S2**. CD9 expression levels in IL-10⁺ versus IL-10⁻ stimulated B cells by condition within stimulation screen experiment.

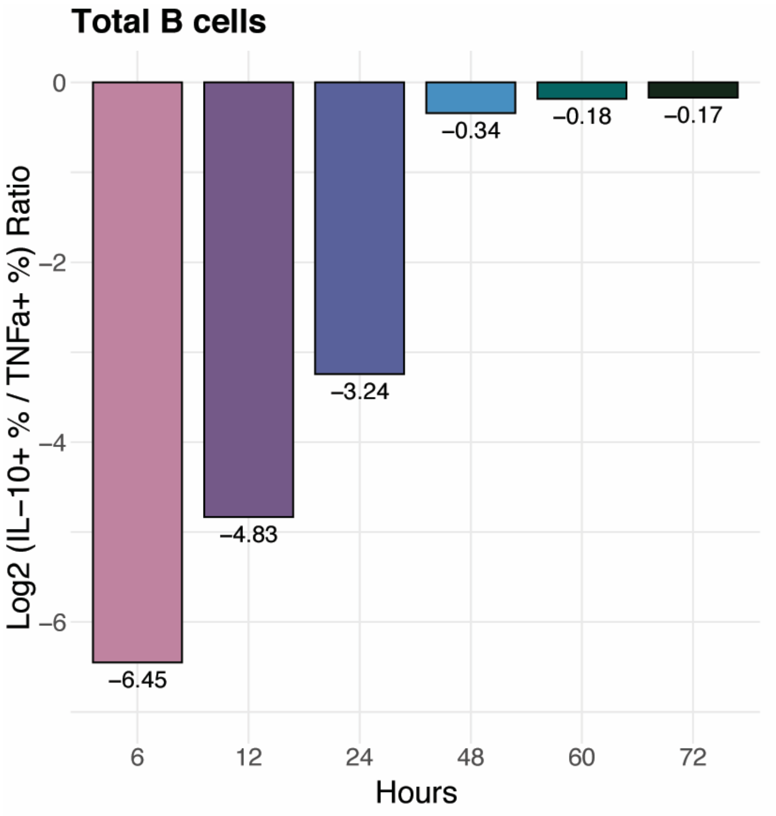


**Figure S3**. Log2 ratio of percentage IL-10$^{+}$ over TNF$\alpha^{+}$ donor-pooled, stimulated B cells by timepoint within timecourse experiment.

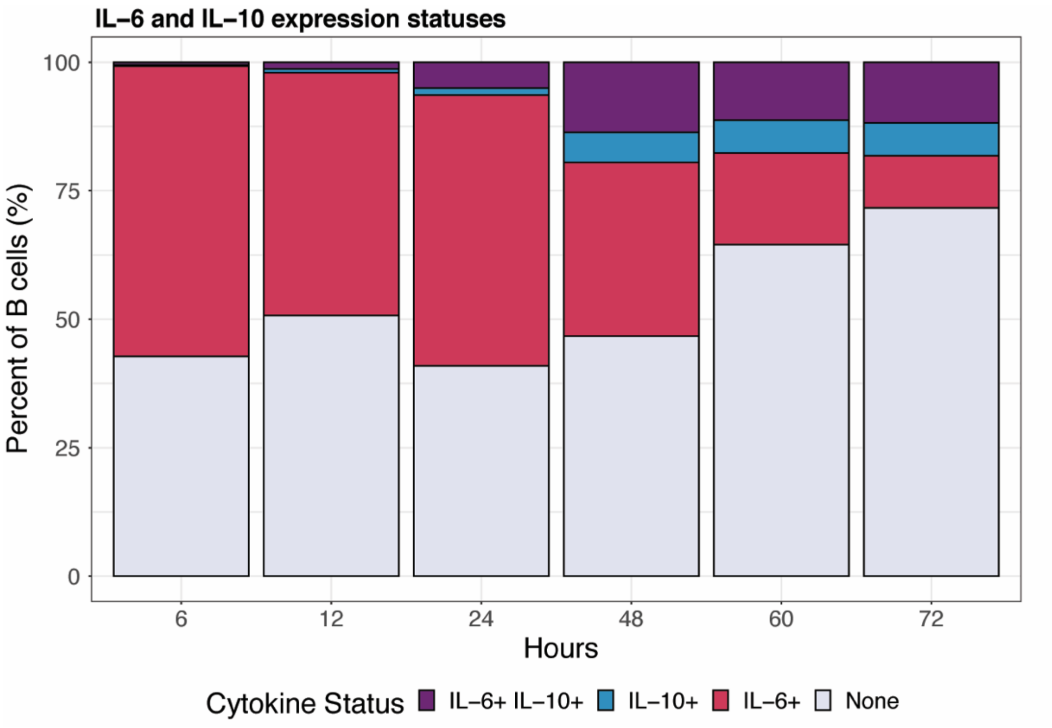


**Figure S4**. Percentage of cells by IL-6 and IL-10 expression for stimulated donor-pooled B cells by timepoint within timecourse experiment.

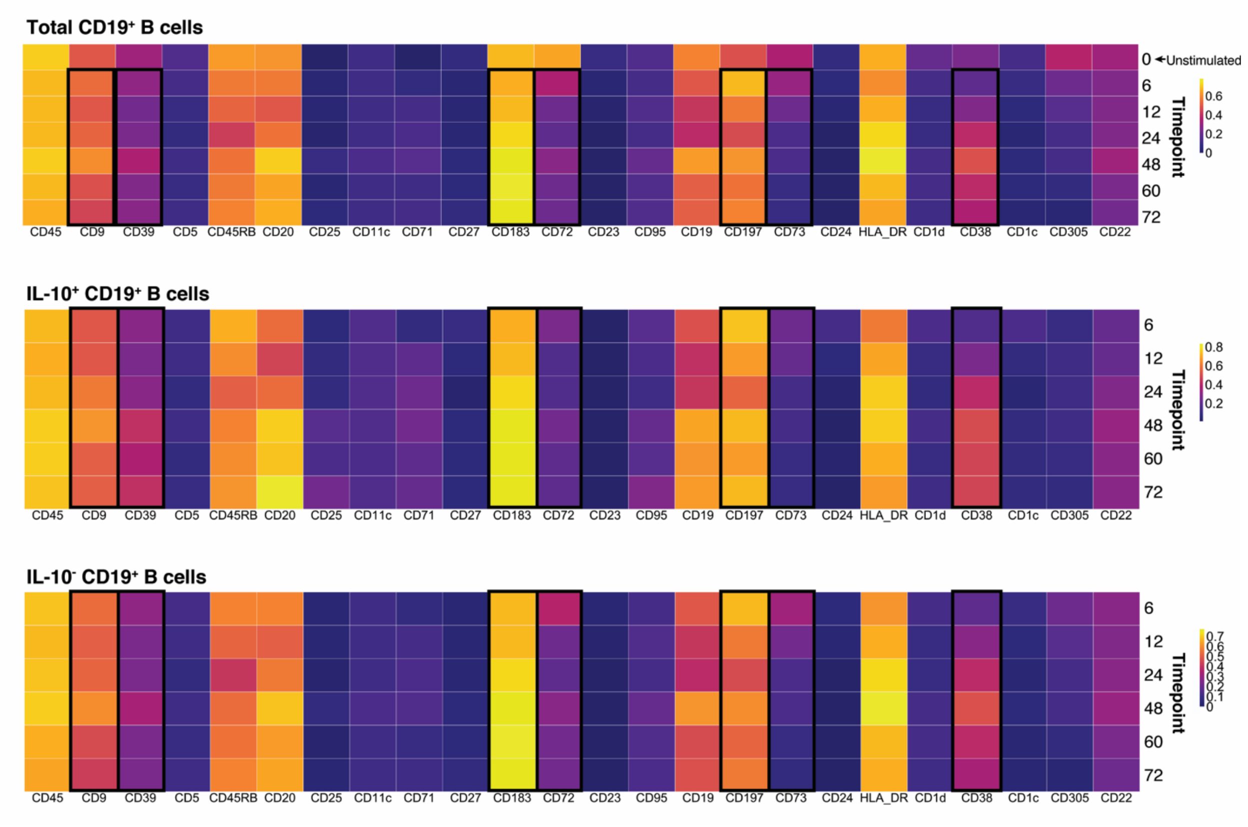


**Figure S5**. Median expression heatmap of total, IL-10$^{+}$ and IL-10$^{-}$ B cells across timepoints of Breg-specific stimulation (6-72 hours).

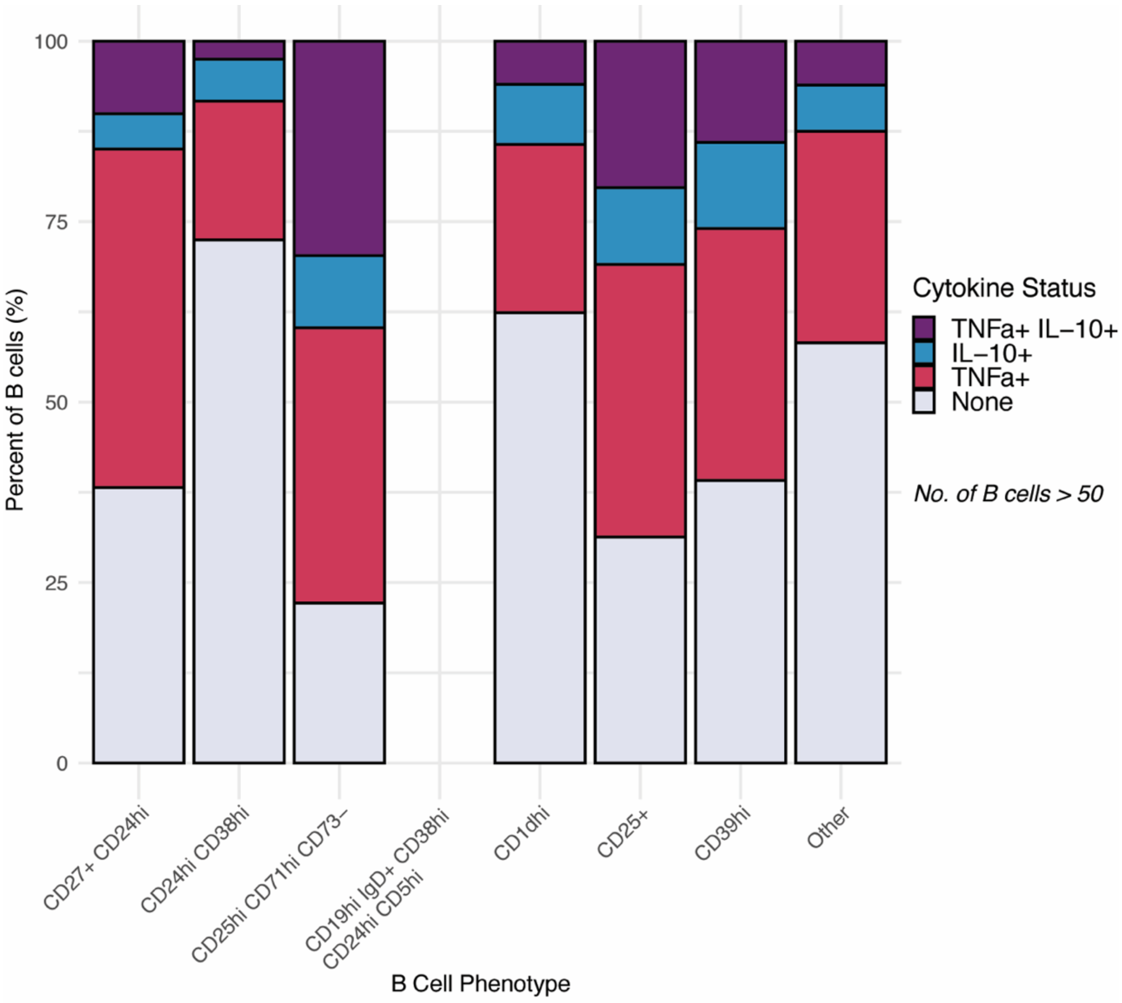


**Figure S6**. Percentage of cells by TNFα and IL-10 expression and B cell phenotype from pooled data of screen and timecourse experiments.

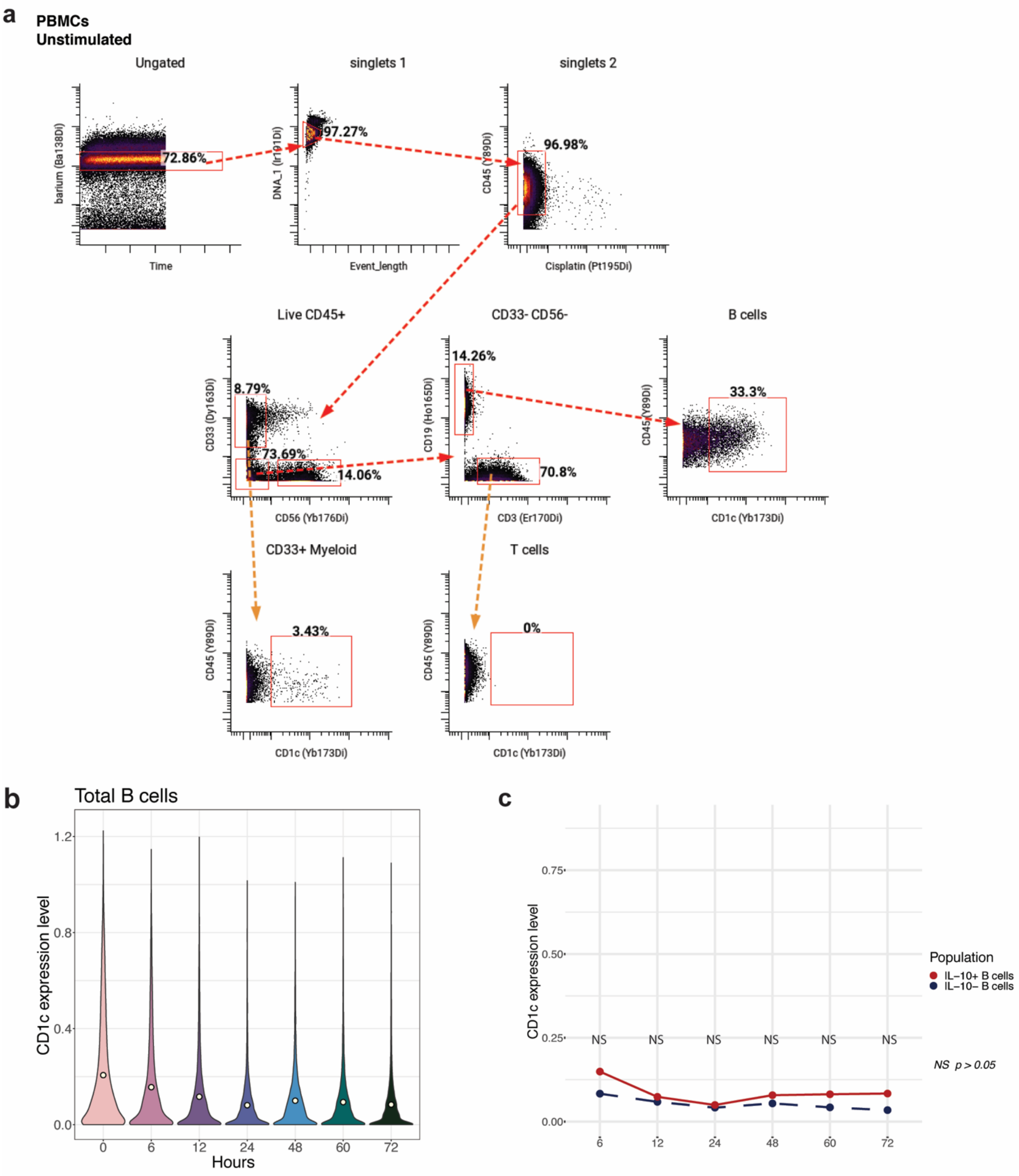


**Figure S7**. Mass cytometry analysis of CD1c expression in unstimulated immune cells and stimulated B cells of healthy individuals.

**(A)** Representative gating strategy for total unstimulated peripheral B cells, T cells and CD33⁺ myeloid cells within mass cytometry analysis of total PBMCs, with exemplary biaxial plots of CD1c expression in each gated cell population.

**(B)** Median CD1c expression levels in total B cells by timepoint in the timecourse experiment. Unstimulated B cell data is depicted as timepoint 0.

**(C)** Median CD1c expression levels in stimulated IL-10⁺ B cells (red) versus IL-10- B cells (blue) by timepoint in the timecourse experiment.

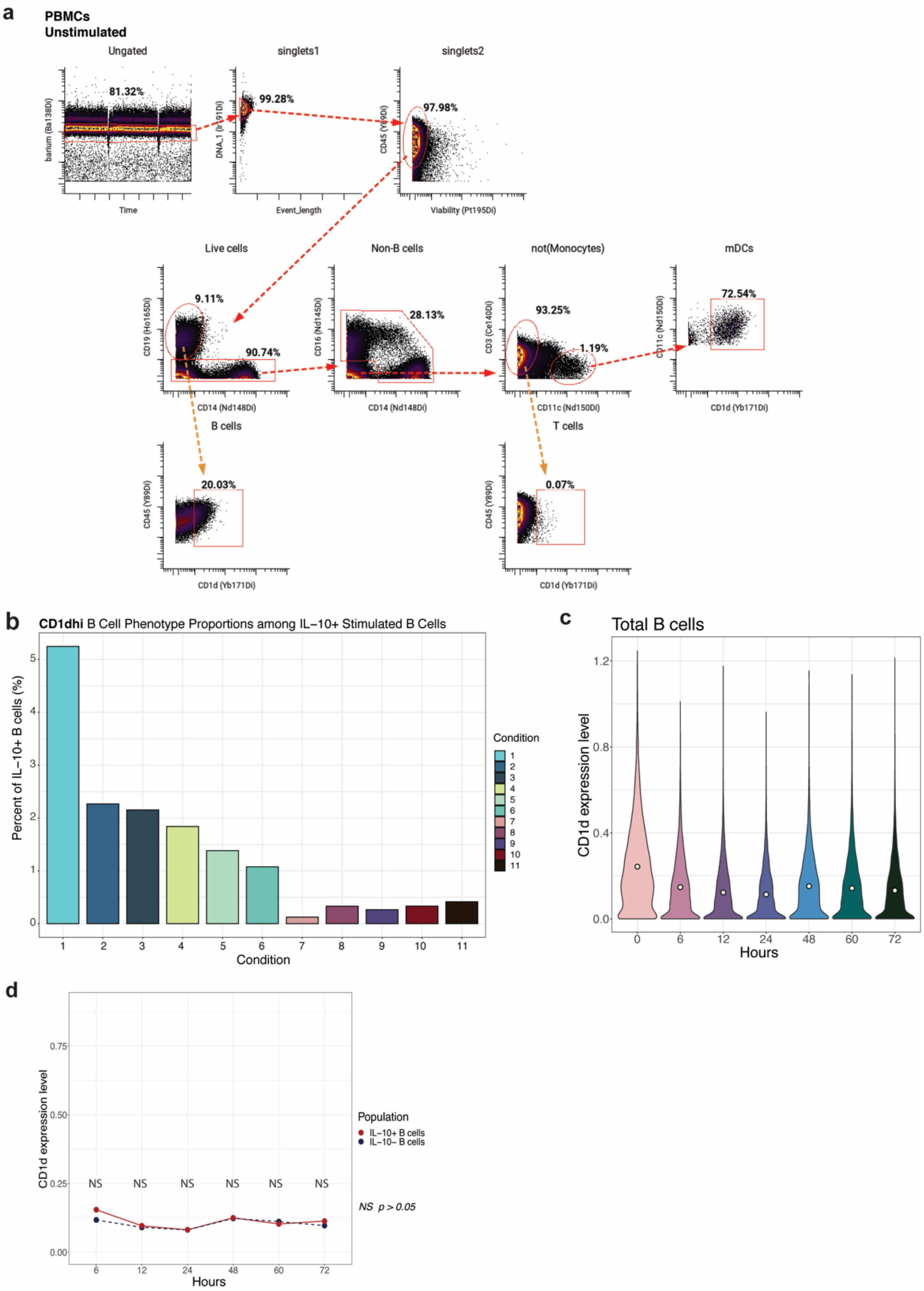


**Figure S8**. Mass cytometry analysis of CD1d expression in total unstimulated immune cells and stimulated B cells of healthy individuals.

**(A)** Representative gating strategy for total unstimulated peripheral B cells, T cells and dendritic cells (DCs) within mass cytometry analysis of total PBMCs, with exemplary biaxial plots of CD1d expression in each gated cell population.

**(B)** Percent of $CD1d^{hi}$ B cells among total IL-$10^{+}$ B cells by stimulation condition in the screen experiment.

**(C)** Median CD1d expression levels in total B cells by timepoint in the timecourse experiment. Unstimulated B cell data is depicted as timepoint 0.

**(D)** Median CD1d expression levels in stimulated IL-$10^{+}$ B cells (red) versus IL-10- B cells (blue) by timepoint in the timecourse experiment.

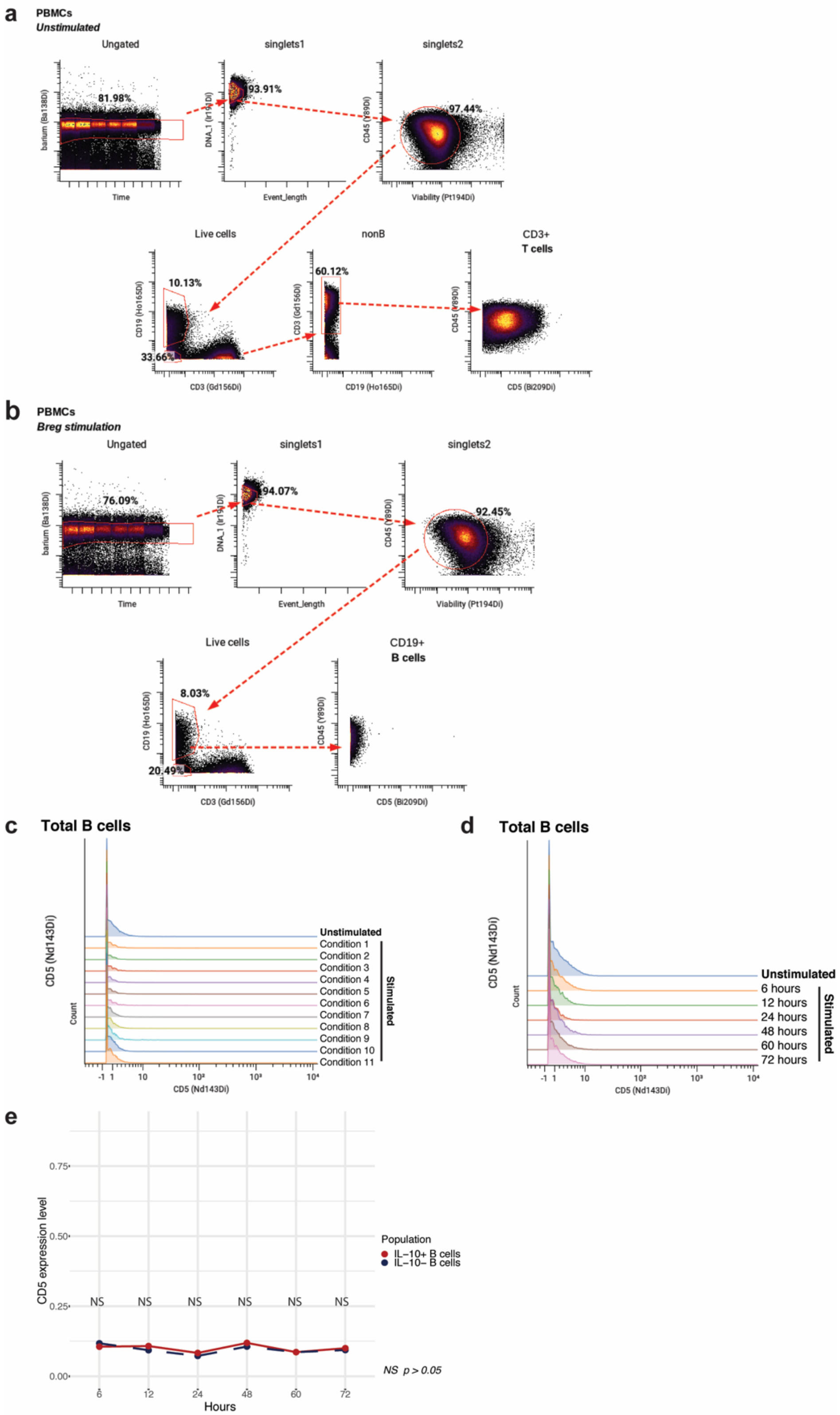


**Figure S9**. Mass cytometry analysis of CD5 expression in unstimulated T and B cells and stimulated B cells of healthy individuals.

**(A)** Representative gating strategy for total unstimulated T cells within mass cytometry analysis of total PBMCs, with final biaxial plot showing positive CD5 antibody staining of T cells.

**(B)** Representative gating strategy for total Breg-stimulated B cells within mass cytometry analysis of total PBMCs, with final biaxial plot indicating minimal CD5 antibody staining of B cells.

**(C)** Overlapping histogram of low or absent CD5 expression level in total B cells in each condition of the Breg stimulation screen.

**(D)** Overlapping histogram of CD5 expression level in total B cells in each experimental timepoint of the Breg stimulation timecourse.

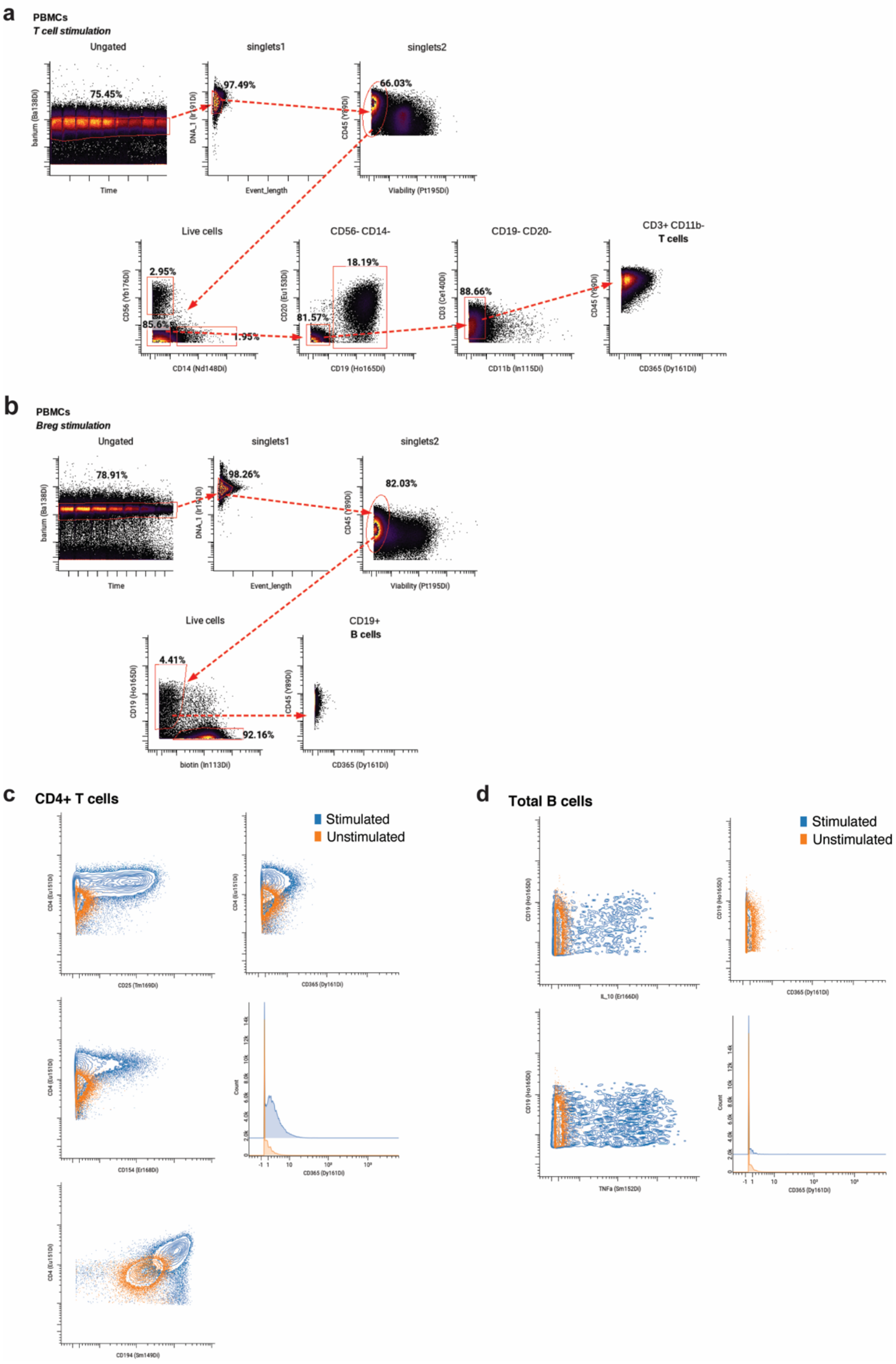


**Figure S10**. Mass cytometry analysis of CD365/TIM-1 expression in unstimulated and stimulated T and B cells of healthy individuals.

**(A)** Representative gating strategy for anti-CD3/anti-CD28-stimulated peripheral T cells within mass cytometry analysis of total PBMCs, with exemplary biaxial plot of CD365/TIM-1 expression in T cells.

**(B)** Representative gating strategy for Breg-stimulated peripheral B cells within mass cytometry analysis of total PBMCs, with exemplary biaxial plot of CD365/TIM-1 expression in B cells.

**(C)** Exemplarily biaxial plots depicting upregulation of activation molecules, CD25, CD154/CD40L and CD194/CCR4, and CD365/TIM-1 expression level in anti-CD3/anti-CD28-stimulated (blue) total peripheral $CD4^+$ T cells as compared to unstimulated (orange) cells from the same healthy individuals.

**(D)** Exemplarily biaxial plots depicting upregulation of the IL-10 and TNF$\alpha$ cytokines, and no change in CD365/TIM-1 expression level in Breg-stimulated (blue) total B cells as compared to unstimulated (orange) cells from the same healthy individuals.

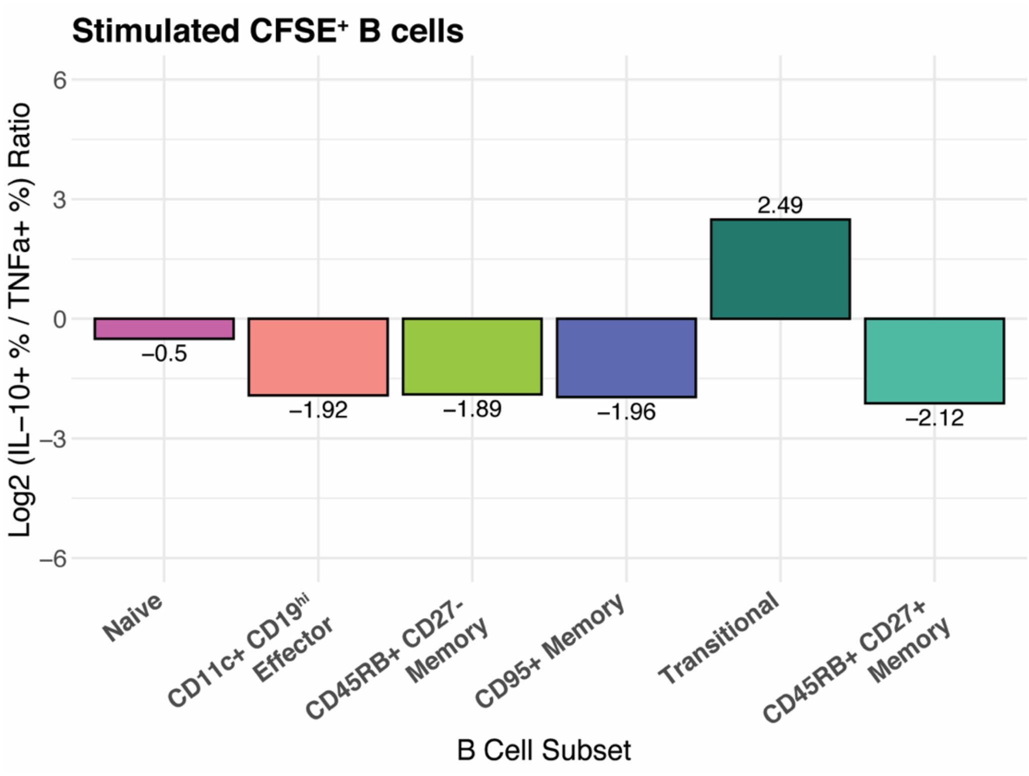


**Figure S11**. Log2 ratio of percentage IL-10$^+$:TNF$\alpha^+$ donor-pooled stimulated B cells by sorted B cell subset.

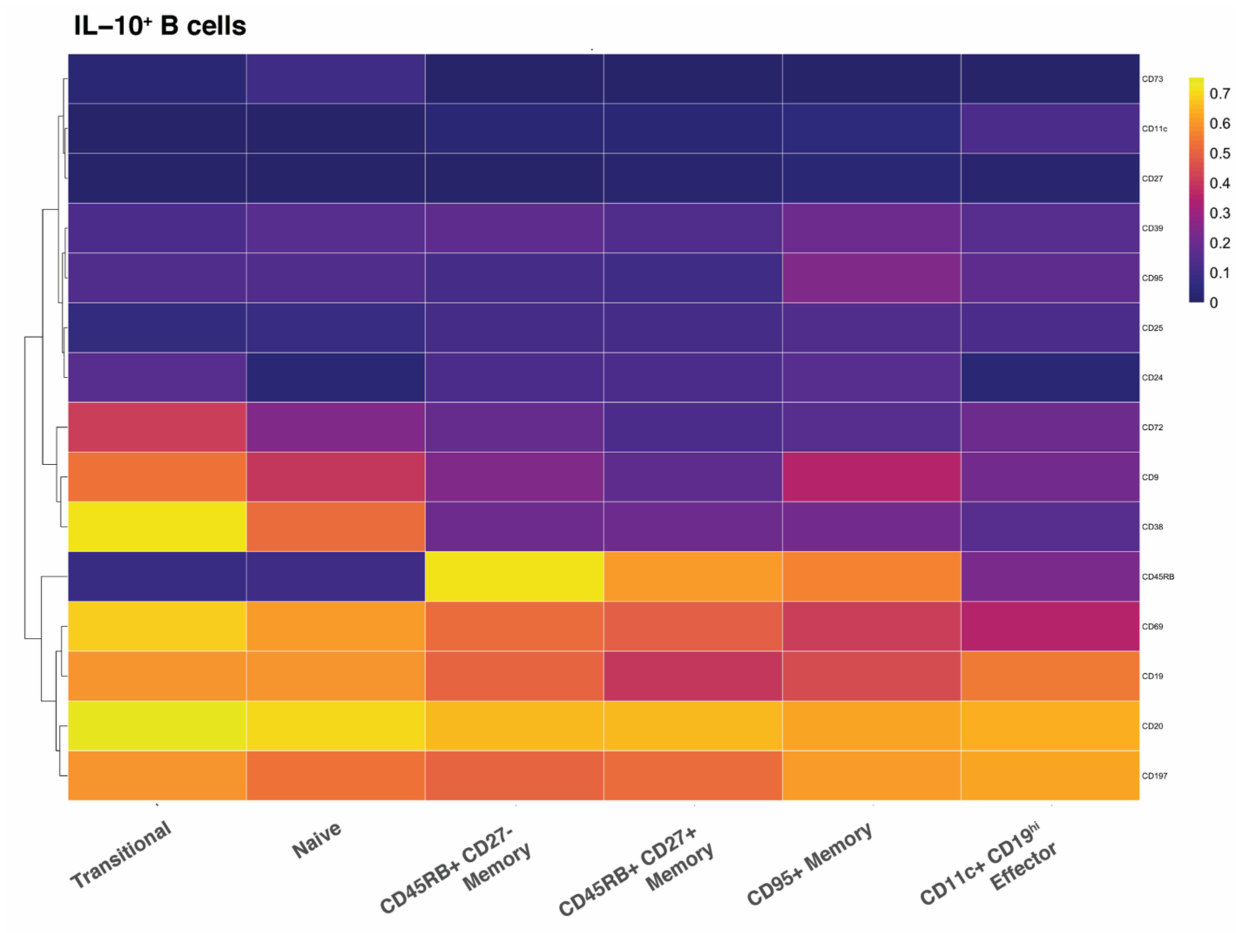


**Figure S12**. Surface marker expression heatmap of IL-10$^{+}$ CFSE$^{+}$ B cell subsets within B cell sorting experiment.

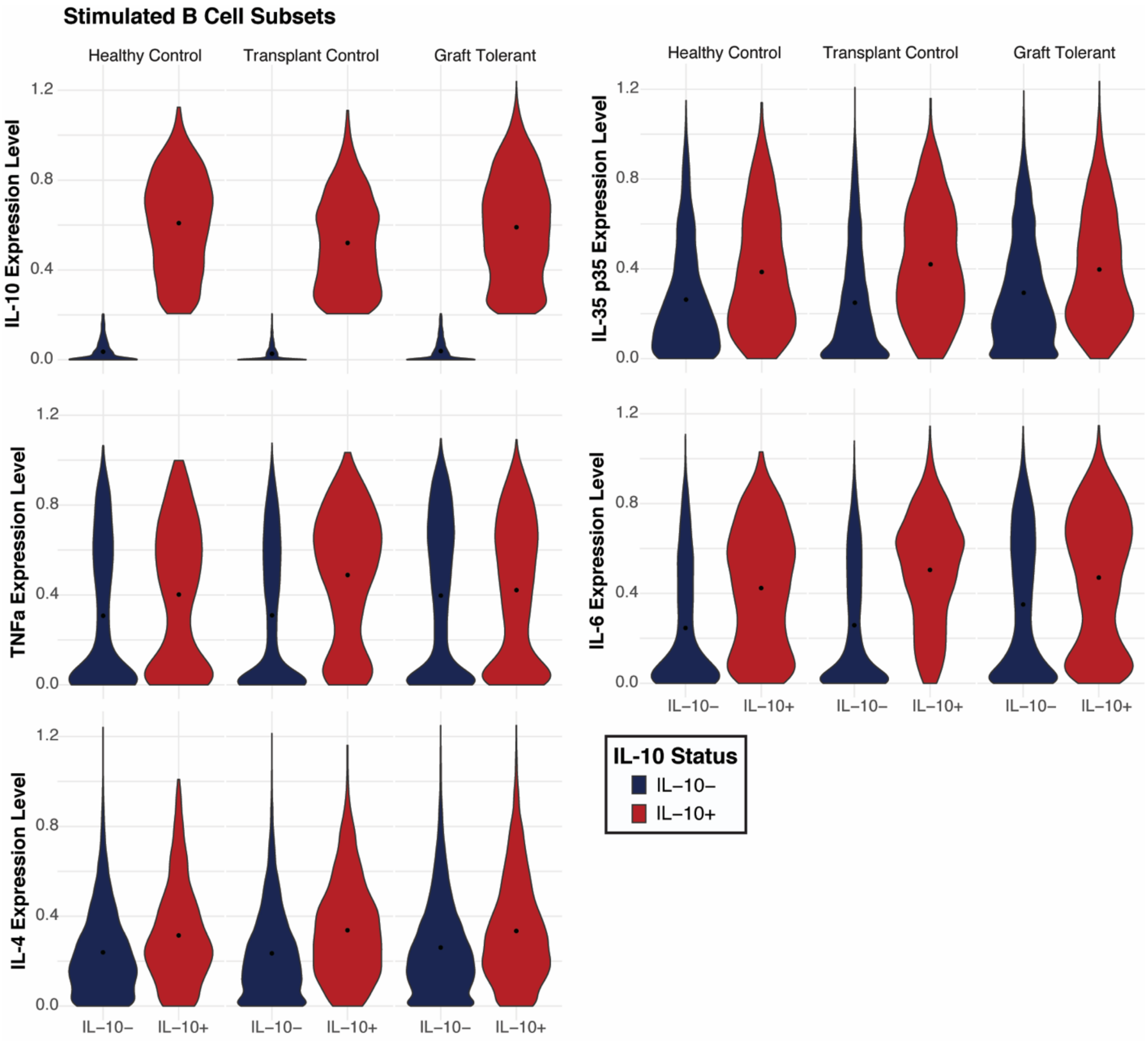


**Figure S13**. Cytokine expression levels of IL-10$^{+}$ and IL-10$^{-}$ stimulated B cells in organ transplant recipient clinical cohorts and healthy control group.

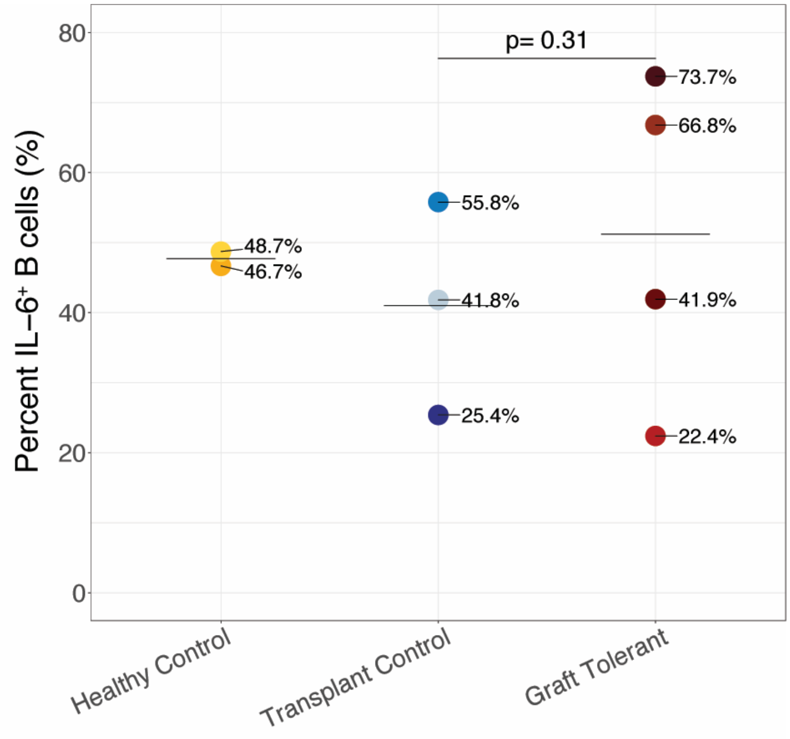


**Figure S14**. Percent of IL-6$^+$ B cells by group and individuals in clinical cohort analysis.

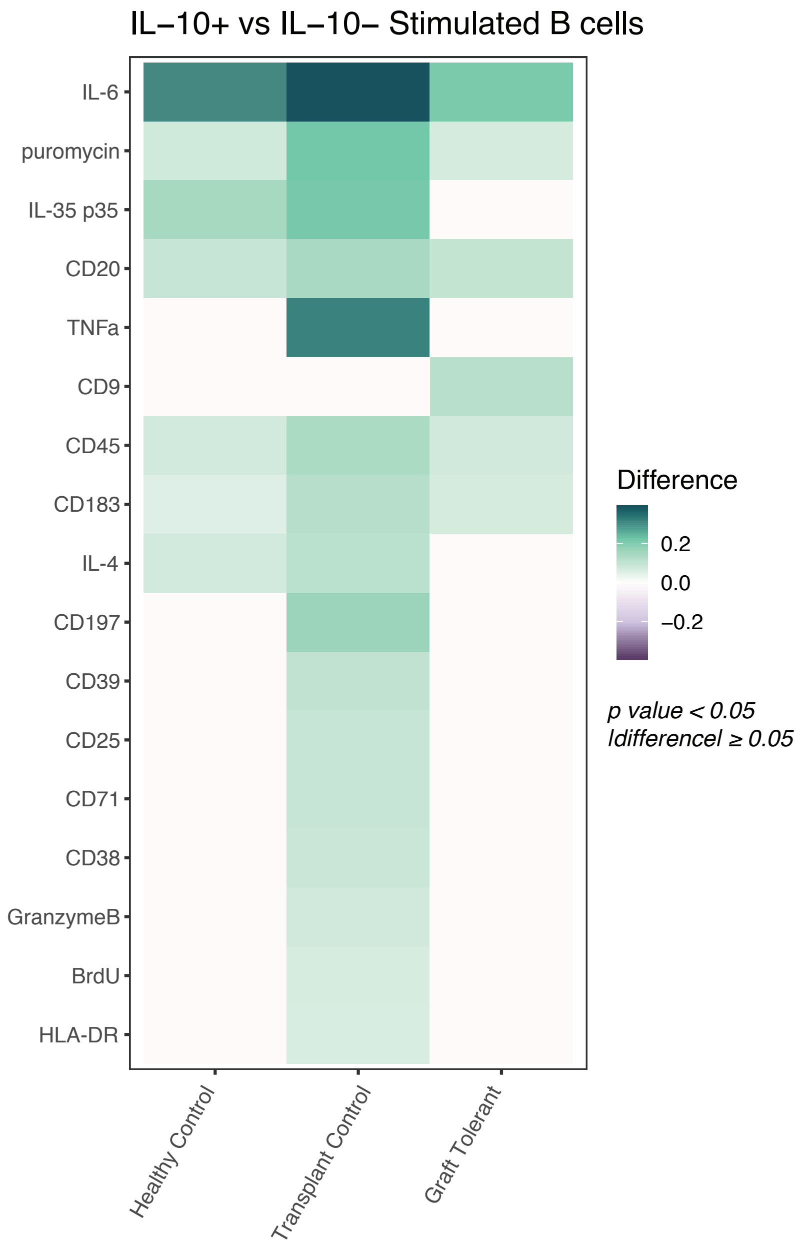


**Figure S15**. Unique expression of surface markers and cytokines in IL-10$^+$ B cells in organ transplant recipient clinical cohorts and healthy control group.

Relative difference in median expression of surface markers and cytokines in IL-10+ versus IL-10- stimulated B cells by clinical cohort. Expression only shown for markers and cytokines with difference greater than or equal to 0.05 and a P value less than 0.05. P values denote result of Kolmogorov-Smirnov test with Bonferroni multiple hypothesis correcting.

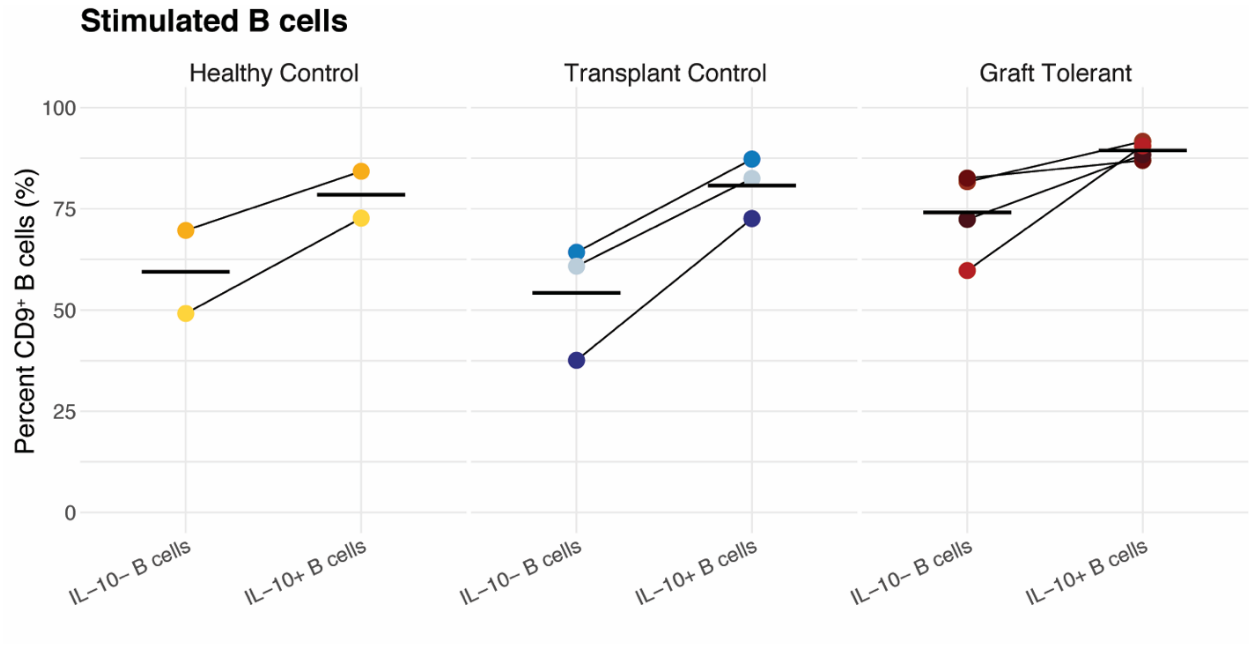


**Figure S16**. Percent of $CD9^{+}$ cells among $IL\text{-}10^{-}$ and $IL\text{-}10^{+}$ stimulated B cells by group and individuals in clinical cohort analysis.

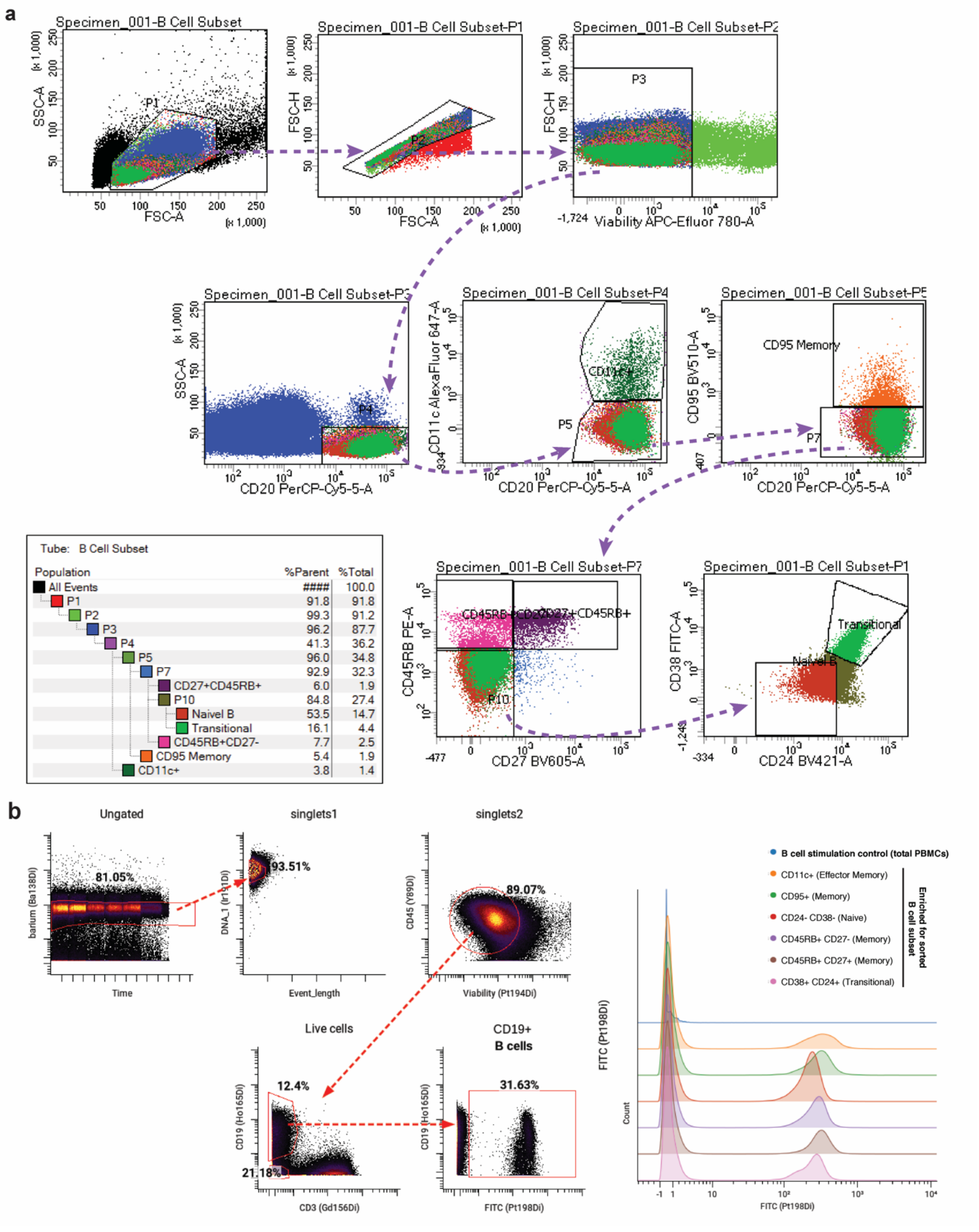


**Figure S17**. Representative gating strategies and CFSE labeling of B cell subsets in sorting experiments.

**(A)** Sorting strategy to isolate CD24$^-$ CD38$^-$ naïve, CD24$^+$ CD38$^+$ transitional, CD45RB$^+$ CD27$^+$ memory, CD45RB$^+$ CD27$^-$ memory, CD95$^+$ memory, and CD11$^+$ effector memory subsets from live CD20$^+$ human peripheral blood B cells for CFSE labeling and *in vitro* Breg stimulation with total PBMCs.

**(B)** Representative gating strategy for total B cells within mass cytometry analysis of B cell subset sorting experiment (left panel). Distinct antibody staining of CFSE$^+$ B cells is indicated in the last biaxial plot of the gating strategy, representing the sorted and CFSE-labeled transitional B cell subset, and in the overlapping histograms representing all sorted and labeled B cell subsets (right panel).

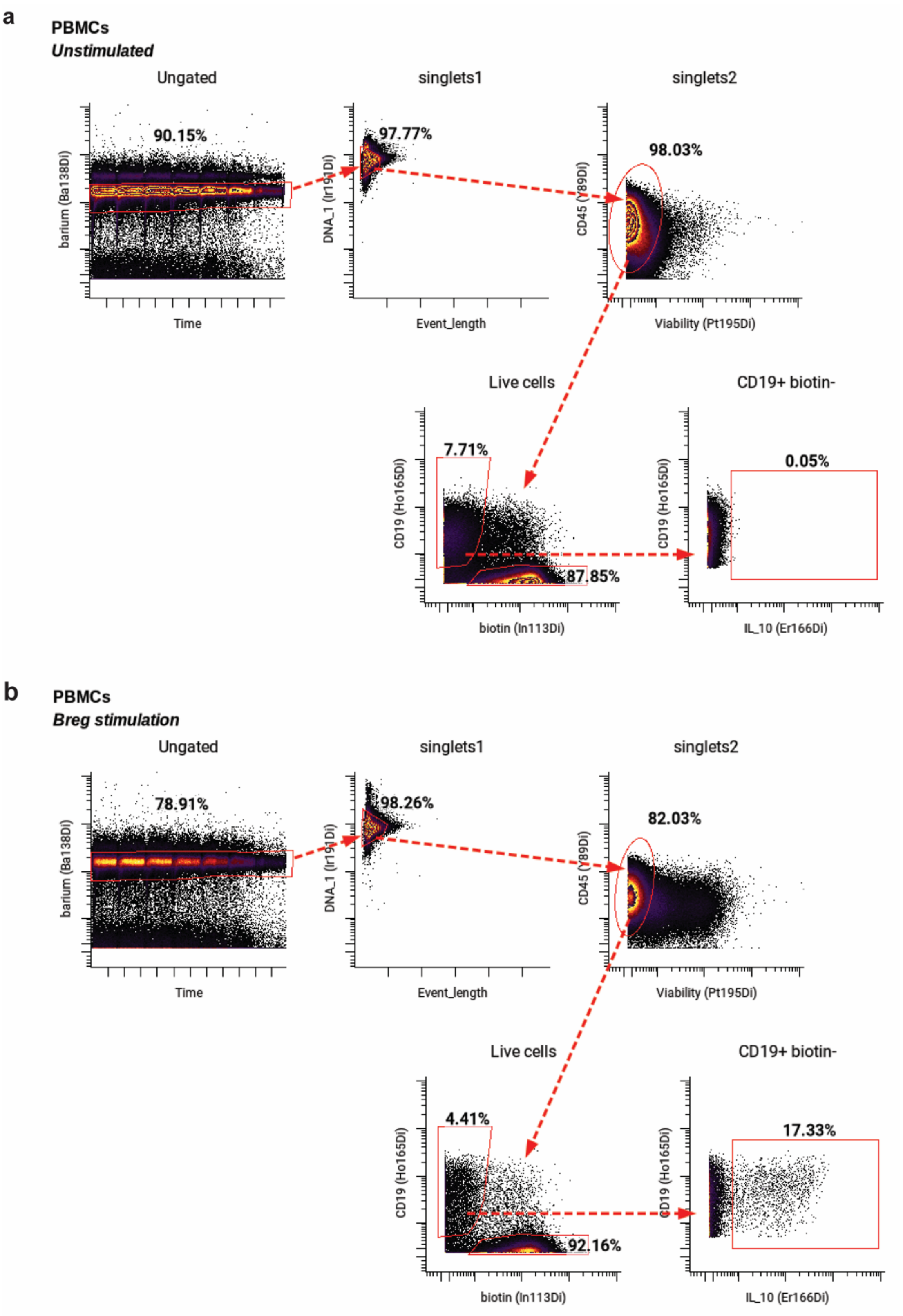


**Figure S18**. Representative gating strategy for unstimulated and stimulated B cells in mass cytometry analysis.

**(A)** Representative gating strategy for total unstimulated peripheral B cells within mass cytometry analysis of total PBMCs, with exemplary biaxial plot indicating absence of IL-10 expression.

**(B)** Representative gating strategy for total stimulated peripheral B cells within mass cytometry analysis of total PBMCs, with exemplary biaxial plot of high IL-10 expression.

**Table S1**. Clinical characteristics of transplant patients and healthy control individuals.

| Clinical group | n | Transplanted Organ(s) | IS at time of sampling? | Rejection episode before or at sampling? | IS | Interval wo IS before sampling range / mean (years) | Interval wo IS after sampling range / mean (years) | Sex ratio (M:F) | Age range (years) | Age mean (years) |
|---|---|---|---|---|---|---|---|---|---|---|
| Operational graft tolerance | 4 | Liver | No | No | ---- | 0.25 - 13 / 7.1* | 0.42 - 5 / 2.8* | 1:1 | 0.42 - 21 | 13.6 |
| Transplant recipient control | 3 | Liver | Yes | No | *Prograf* | ---- | ---- | 1:1 | 7 - 19 | 14.8 |
| Healthy control | 2 | None | No | ---- | ---- | ---- | ---- | 1:1 | 19 - 32 | 25.5 |

** Data not available for one individual*

Abbreviations: *IS, Immunosuppression*

**Table S2**. Frequency of live cells and total and IL-10$^+$ B cells in clinical groups and healthy individuals.

| Donor | Status | Condition | Proportion live CD45+ cells among singlets (%) | Proportion B cells among live CD45+ cells (%) | Proportion IL-10+ B cells among total B cells (%) |
|---|---|---|---|---|---|
| HCON1 | Healthy Control | Unstimulated | 98.03 | 8.05 | NA |
| HCON1 | Healthy Control | Stimulated | 72.05 | 1.33 | 25.42 |
| HCON2 | Healthy Control | Unstimulated | 98.61 | 7.62 | NA |
| HCON2 | Healthy Control | Stimulated | 81.82 | 1.37 | 28.89 |
| TCON1 | Transplant Control | Unstimulated | 97.40 | 7.97 | NA |
| TCON1 | Transplant Control | Stimulated | 84.32 | 4.41 | 8.09 |
| TCON2 | Transplant Control | Unstimulated | 39.41 | 1.38 | NA |
| TCON2 | Transplant Control | Stimulated | 66.76 | 2.88 | 2.57 |
| TCON3 | Transplant Control | Unstimulated | 86.44 | 7.15 | NA |
| TCON3 | Transplant Control | Stimulated | 90.65 | 2.02 | 9.31 |
| TOL1 | Graft Tolerant | Unstimulated | 79.91 | 17.28 | NA |
| TOL1 | Graft Tolerant | Stimulated | 70.63 | 12.23 | 23.70 |
| TOL2 | Graft Tolerant | Unstimulated | 67.29 | 5.50 | NA |
| TOL2 | Graft Tolerant | Stimulated | 57.81 | 3.14 | 24.43 |
| TOL3 | Graft Tolerant | Unstimulated | 67.22 | 6.84 | NA |
| TOL3 | Graft Tolerant | Stimulated | 82.03 | 4.64 | 16.74 |
| TOL4 | Graft Tolerant | Unstimulated | 95.20 | 7.69 | NA |
| TOL4 | Graft Tolerant | Stimulated | 36.76 | 8.69 | 28.03 |

**Table S3**. Reagents for PBMC *in vitro* stimulations.

| Experiment | Condition | Reagent | Final concentration | Vendor | Catalogue number |
|---|---|---|---|---|---|
| Cytokine and phenotypic screen of B cell stimulations (Fig 1 and 3) | 1 - 3 | R848 | 1 ug/mL | Mabtech | 3660-1 |
| | 4 - 6 | Lipopolysaccharides (LPS) from Escherichia coli O127:B8 | 10 ug/mL | Sigma | L4516-1MG |
| | 7 - 11 | CpG ODN 2006 | 10 ug/mL | Invivogen | tlrl-2006 |
| | | Soluble human anti-CD40 clone 82111 | 500 ng/mL | RD Systems | MAB6321-100 |
| | 3, 6 | Recombinant human IL-4 | 20 ng/mL | RD Systems | 204-IL-020 |
| | 3, 6, 9, 11 | Recombinant human IL-10 | 25 ng/mL | Biolegend | 571002 |
| | 9 - 11 | Recombinant human IL-2 | 600 IU/mL | Mabtech | 3660-1 |
| | 9 - 10 | Recombinant human IL-21 | 100 ng/mL | RD Systems | 8879-IL-010 |
| | 10 - 11 | Recombinant human IL-35 | 20 ng/mL | Enzo Life | ALX-522-140-C010 |
| | All | Phorbol 12-myristate 13-acetate | 50 ng/mL | Sigma | P8139-1MG |
| | All | Ionomycin calcium salt from Streptomyces conglobatus | 1 ug/mL | Sigma | I0634-1MG |
| | All | Brefeldin A Solution | 1X | eBioscience | 00-4506-51 |
| Timecourse of Breg stimulation (Fig 2-3), B cell subset sorting and stimulation (Fig 4), and clnical cohort and control B cell analysis (Fig 5) | All | CpG ODN 2006 | 10 ug/mL | Invivogen | tlrl-2006 |
| | All | Soluble human anti-CD40 clone 82111 | 500 ng/mL | RD Systems | MAB6321-100 |
| | All | Recombinant human IL-2 | 600 IU/mL | Mabtech | 3660-1 |
| | All | Recombinant human IL-21 | 100 ng/mL | RD Systems | 8879-IL-010 |
| | All | Recombinant human IL-35 | 20 ng/mL | Enzo Life | ALX-522-140-C010 |
| | All | Phorbol 12-myristate 13-acetate | 50 ng/mL | Sigma | P8139-1MG |
| | All | Ionomycin calcium salt from Streptomyces conglobatus | 1 ug/mL | Sigma | I0634-1MG |
| | All | Brefeldin A Solution | 1x | eBioscience | 00-4506-51 |
| T cell stimulation to validate antibody staining (Fig S...) | All | Plate-bound human anti-CD3 | 10 ug/mL | Biolegend | 302933 |
| | All | Plate-bound human anti-CD28 | 5 ug/mL | Biolegend | 317325 |
| | All | Recombinant human IL-4 | 12.5 ng/mL | RD Systems | 204-IL-020 |
| | All | Recombinant human IL-2 | 50 IU/mL | Biolegend | 589102 |
| | All | Brefeldin A Solution | 1x | eBioscience | 00-4506-51 |

**Table S4**. Antibody panel used for mass cytometry analysis.

| Experiment | Antigen | Clone | Tag | Vendor |
|---|---|---|---|---|
| **Panel 1:** Mass cytometry analysis of healthy human peripheral B cells (Fig 1-3) and organ transplant recipient peripheral B cells (Fig 5) | CD45 | HI30 | 89 Y | Biolegend |
| | CD9 | HI9a | 141 Pr | Biolegend |
| | CD39/ENTPD1 | A1 | 142 Nd | Biolegend |
| | CD5 | UCHT2 | 143 Nd | Biolegend |
| | CD45RB | MEM-55 | 145 Nd | Fluidigm |
| | CD20 | 2H7 | 147 Sm | Fluidigm |
| | CD25 | BC96 | 149 Sm | Biolegend |
| | CD11c | Bu15 | 150 Nd | Biolegend |
| | CD71 | CY1G4 | 151 Eu | Biolegend |
| | IgK | A8B5 | 153 Eu | Invitrogen |
| | CD27 | M-T271 | 155 Gd | Biolegend |
| | CD183 | G025H7 | 156 Gd | Fluidigm |
| | CD72 | 3F3 | 157 Gd | Biolegend |
| | CD23 | EBVCS-5 | 160 Gd | Biolegend |
| | CD365/TIM-1 | 1D12 | 161 Dy | Biolegend |
| | CD95/FASR | DX2 | 164 Dy | Biolegend |
| | CD19 | HIB19 | 165 Ho | Biolegend |
| | CD197/CCR7 | G043H7 | 167 Er | Fluidigm |
| | CD73/NT5E1 | AD2 | 168 Er | Biolegend |
| | CD24 | ML5 | 169 Tm | Biolegend |
| | HLA-DR | L243 | 170 Er | Fluidigm |
| | CD1d | 51.1 | 171 Yb | Biolegend |
| | CD38 | HIT2 | 172 Yb | Biolegend |
| | CD1c | L161 | 173 Yb | Biolegend |
| | IgL | MHL-38 | 174 Yb | BD Biosciences |
| | CD305/LAIR1 | NKTA255 | 176 Yb | Genetex |
| | CD22 | HIB22 | 209 Bi | Biolegend |
| | Biotin | 1D4-C5 | 113 In | Biolegend |
| | IgM | polyclonal | 140 Ce | Invitrogen |
| | IL-4 | MP4-25D2 | 144 Nd | Fluidigm |
| | IgD | IA6-2 | 146 Nd | Biolegend |
| | IgA | polyclonal | 148 Nd | Fluidigm |
| | TNFa | Mab11 | 152 Sm | Fluidigm |
| | IL-6 | MQ2-13A5 | 154 Sm | Biolegend |
| | Puromycin | 12D10 | 158 Gd | Miltenyi Biotec |
| | IL-35/IL-12 p35 | 27537 | 159 Tb | RD Systems |
| | IgG | M1310G05 | 162 Dy | Biolegend |
| | Granzyme B | 351927 | 163 Dy | RD Systems |
| | IL-10 | JES3-9D7 | 166 Er | Biolegend |
| | BrU | 3D4 | 175 Lu | BD Biosciences |
| | CD3 | OKT3 | Biotin | Biolegend |
| | CD7 | CD7-6B6 | Biotin | Miltenyi Biotec |
| | CD15 | HI98 | Biotin | Biolegend |
| | CD33 | WM53 | Biotin | Biolegend |
| | CD56 | 5.1H11 | Biotin | Biolegend |
| | CD61 | Y2/51 | Biotin | Miltenyi Biotec |
| | CD235ab | HIR2 | Biotin | Biolegend |
| **Panel 2:** Sorting B cell subsets (Fig 4) | CD20 | 2H7 | PerCP-Cy5.5 | Biolegend |
| | CD38 | HIT2 | FITC | Biolegend |
| | CD24 | ML5 | Brilliant Violet 421 | Biolegend |
| | CD45RB | MEM-55 | PE | Biolegend |
| | CD27 | O323 | Brilliant Violet 605 | Biolegend |
| | CD95 | DX2 | Brilliant Violet 510 | Biolegend |
| | CD11c | Bu15 | Alexa Fluor 647 | Biolegend |

| | | | | |
|---|---|---|---|---|
| **Panel 3**: Mass cytometry analysis of healthy donor peripheral B cells in sorting experiment (Fig 4) | CD45 | HI30 | 89 Y | Biolegend |
| | CD14 | M5E2 | 113 In | Biolegend |
| | CD11b | ICRF44 | 115 In | Biolegend |
| | CD9 | HI9a | 141 Pr | Biolegend |
| | CD39/ENTPD1 | A1 | 142 Nd | Biolegend |
| | CD16 | 3G8 | 143 Nd | Biolegend |
| | CD69 | FN50 | 144 Nd | Fluidigm |
| | CD45RB | MEM-55 | 145 Nd | Fluidigm |
| | CD20 | 2H7 | 147 Sm | Fluidigm |
| | CD25 | BC96 | 149 Sm | Biolegend |
| | CD11c | Bu15 | 150 Nd | Biolegend |
| | CD4 | RPA-T4 | 151 Eu | Biolegend |
| | TIGIT | MAB7898 | 153 Eu | Fluidigm |
| | CD27 | M-T271 | 155 Gd | Biolegend |
| | CD3 | UCHT1 | 156 Gd | Biolegend |
| | CD72 | 3F3 | 157 Gd | Biolegend |
| | CD95/FASR | DX2 | 164 Dy | Biolegend |
| | CD19 | HIB19 | 165 Ho | Biolegend |
| | CD197/CCR7 | G043H7 | 167 Er | Fluidigm |
| | CD24 | ML5 | 169 Tm | Biolegend |
| | HLA-DR | L243 | 170 Er | Fluidigm |
| | CD8a | RPA-T8 | 171 Yb | Biolegend |
| | CD38 | HIT2 | 172 Yb | Biolegend |
| | CD73/NT5E | AD2 | 173 Yb | Biolegend |
| | CD279/PD1 | EH12.2H7 | 174 Yb | Fluidigm |
| | CD127 | A019D5 | 176 Yb | Biolegend |
| | CD5 | UCHT2 | 209 Bi | Biolegend |
| | IgM | polyclonal | 140 Ce | Invitrogen |
| | IgD | IA6-2 | 146 Nd | Biolegend |
| | IgA | polyclonal | 148 Nd | Fluidigm |
| | TNFa | Mab11 | 152 Sm | Fluidigm |
| | IL-6 | MQ2-13A5 | 154 Sm | Biolegend |
| | Puromycin | 12D10 | 158 Gd | Miltenyi Biotec |
| | IL35/IL12 p35 | 27537 | 159 Tb | RD Systems |
| | CD152/CTLA4 | 14D3 | 160 Gd | Invitrogen |
| | IL17A | BL168 | 161 Dy | Fluidigm |
| | IgG | M1310G05 | 162 Dy | Biolegend |
| | IL-10 | JES3-9D7 | 166 Er | Biolegend |
| | IFNg | B27 | 168 Er | Fluidigm |
| | BrU | 3D4 | 175 Lu | BD Biosciences |
| | CFSE/FITC | polyclonal | 198 Pt | Biolegend |
